# Circadian reprogramming of adipose progenitor cells regulates intermittent fasting-mediated adipose tissue remodeling and metabolic improvement

**DOI:** 10.1101/2022.12.10.519916

**Authors:** Ju Hee Lee, Yash Patel, Joanna Lan-Hing Yeung, Lauren Pickel, Kafi N. Ealey, Jacques Togo, Yun Hye Kim, Kyoung-Han Kim, Jin-Gyoon Park, Timothy Jackson, Allan Okrainec, Jae-Ryong Kim, So-Young Park, Satya Dash, Hoon-Ki Sung

**Author notes:** **Correspondence** should be addressed to Hoon-Ki Sung (H.-K.S.).

## Abstract

White adipose tissue (WAT) fibrosis is a hallmark of dysfunctional WAT that is directly linked to metabolic abnormalities. Recent studies have highlighted the role of dysfunctional adipose progenitor cells (APCs) in WAT fibrosis and impaired adaptive tissue plasticity, leading to systemic insulin resistance. However, therapeutic options for WAT fibrosis are limited. Intermittent fasting (IF) is an effective dietary regimen for weight control and metabolic improvement through various mechanisms, including healthy remodeling of WAT. However, whether IF is effective in improving age-associated WAT fibrosis and metabolic homeostasis is unknown. Here, we show that IF confers therapeutic benefits in aged and obese mice through reduction of WAT fibrosis. Single-cell analyses revealed that IF significantly reduces pro-fibrotic signatures within APCs along with upregulation of the circadian pathways, suggesting that the circadian clock of APCs mediates IF-induced WAT remodeling. Importantly, mice lacking core circadian gene exhibited increased fibrotic signatures in WAT and diminished beneficial response to IF, further supporting the importance of circadian rhythm in IF-mediated metabolic benefits. Lastly, insulin resistance in humans also presented with dysregulated circadian rhythm signatures in APC populations. Collectively, our findings highlight the novel role of the APC circadian rhythm in plasticity of WAT and metabolic response to IF.

## INTRODUCTION

The white adipose tissue (WAT) is a metabolically active endocrine organ that plays a pivotal role in whole-body energy metabolism through dynamic remodeling in response to environmental cues and energy balance. However, the remodeling capacity of WAT is significantly attenuated with ageing and chronic overnutrition, which drive pathological adipose remodeling. Adipose fibrosis is a hallmark of dysfunctional WAT (Pellegrinelli et al., 2016; Rutkowski et al., 2015). Importantly, WAT fibrosis is directly linked to metabolic disease and insulin resistance (Hasegawa et al., 2018; Palmer and Kirkland, 2016; Sun et al., 2014; Sun et al., 2013). Recent studies have demonstrated adipose progenitor cells (APCs) to be a major source of extracellular matrix (ECM) production in WAT (Marcelin *et al*., 2017; Pellegrinelli *et al*., 2016). Under obesogenic conditions, a subset of APCs adopt a pro-fibrotic phenotype characterized by high expression of CD9 (CD9^high^) and play a key role in WAT fibrosis and impairment of adaptive tissue plasticity (Marcelin *et al*., 2017). In contrast, an adipogenic CD9^low^ subset of APCs is largely diminished in obese WAT, resulting in unhealthy WAT expansion via adipocyte hypertrophy instead of healthy hyperplastic expansion (Pellegrinelli *et al*., 2016; Rutkowski *et al*., 2015). Similarly, low-adipogenic, high fibro-inflammatory progenitors, Ly6C+/PDGFRβ+ APCs, accumulate in the WAT of high fat diet (HFD)-fed animals (Hepler et al., 2018). Moreover, the APCs of aged mice were shown to express high levels of senescence signatures and were directly associated with the inability to induce *de novo* beige adipogenesis under cold exposure (Berry et al., 2017). *In vitro* studies also demonstrate that preadipocytes in close contact with inflammatory macrophages, are major producers of pro-fibrotic molecules (e.g., inhibin βA) and ECM components (e.g., collagen and fibronectin), driving increased inflammatory remodeling (Keophiphath et al., 2009). Senescent APCs from human adipose tissue impair the adipogenic potential of healthy co-cultured adipose progenitors, further aggravating the unhealthy microenvironment and reducing tissue plasticity (Xu et al., 2015). Taken together, these phenotypic changes in APCs not only affect their adipogenic and thermogenic capacity, but also play a key role in the fibro-inflammation and therefore metabolic function of the WAT as a whole.

To combat obesity and its associated metabolic abnormalities, dietary regimens, such as intermittent fasting (IF) and time-restricted feeding (TRF), have been extensively investigated. Many studies collectively report that IF confers system-wide effects, including body weight and fat mass reduction, improved glucose/insulin homeostasis, gut microbiome changes, and autophagy activation in metabolic organs (Beli et al., 2018; Cignarella et al., 2018; Kim et al., 2017; Li et al., 2017; Martinez-Lopez et al., 2017). IF has been shown to exert anti-inflammatory and pro-angiogenic effects on WAT, mediating thermogenic activation (browning) and protection against diet-induced obesity (Kim *et al*., 2017; Li *et al*., 2017). Similarly, numerous studies have reported that TRF protects animals from diet-induced obesity by enhancing energy expenditure and preventing metabolic abnormalities such as fatty liver and insulin resistance, in the absence of a reduction in caloric intake (Chaix et al., 2021; Chaix et al., 2019; Hatori et al., 2012). This suggests that incorporating prolonged fasting confers metabolic benefits, regardless of caloric intake reduction. However, although many studies have explored the preventive effects of IF, whether IF can exert therapeutic benefits in animals with established severe metabolic dysfunction is unclear.

Circadian clock genes modulate the temporal expression and function of metabolic organs, optimizing energy metabolism in relation to the daily cycles of food intake and energy expenditure. Circadian rhythm disruption plays a crucial role in the pathogenesis of metabolic diseases and organ dysfunction. Obesogenic diets promptly disrupt the diurnal rhythm in feeding behaviour and alter the expression and cycling of key circadian clock genes and downstream metabolic regulators in various tissues, such as the liver, hypothalamus, and adipose tissue (Kohsaka et al., 2007). Mice deficient in *Bmal1*, a core clock gene, had a complete loss of rhythmic cycling of circadian genes, and they exhibited premature age-related pathologies as well as dysregulated glucose and insulin homeostasis (Kondratov et al., 2006; Lee et al., 2011; Marcheva et al., 2010). Deletion of adipocyte-*Bmal1* has been shown to result in obesogenic phenotypes, including weight gain, adipocyte hypertrophy, and dampened energy expenditure (Paschos et al., 2012).

WAT is a highly rhythmic organ, and various cell types, such as adipocytes and APCs, exhibit rhythmic expression of circadian genes (Man et al., 2020; Pickel and Sung, 2020; Ribas-Latre et al., 2021). The circadian clock-regulated diurnal pattern of *in vivo* APC proliferation was altered by HFD to drive the hyperplastic expansion of WAT (Ribas-Latre *et al*., 2021). The expressions of core circadian genes have been shown to be significantly altered in both adipocytes and APCs in obese human fat, resulting in the activation of numerous target genes involved in WAT inflammation (Maury et al., 2021). Intriguingly, abnormal circadian rhythm was also associated with fibrosis of the liver and lungs, indicating the importance of circadian rhythm in tissue homeostasis and function (Cunningham et al., 2020; Louis et al., 2022; Pekovic-Vaughan et al., 2014; Rainer, 2020). As the expression of circadian genes has been significantly changed in APCs in obese fat (Maury *et al*., 2021), and APCs have been shown to be a key player in WAT fibrosis (Marcelin *et al*., 2017), it is plausible that the abnormal rhythmicity of APCs may be linked to WAT fibrosis and dysfunction. However, the relationship between the circadian dysregulation of APCs and WAT fibrosis is poorly understood. Moreover, since feeding and fasting are the potent cues (zeitgebers) for WAT clocks (Pickel and Sung, 2020), feeding pattern changes can directly affect (entrain) the circadian rhythms of WAT. In this study, we explored the impact of IF on WAT in an aged- and diet-induced obese mouse model. We found that IF alleviates WAT fibrosis and improves insulin resistance through the enhanced circadian rhythm of APCs. These effects were largely diminished in mice lacking the core clock gene *Bmal1*, suggesting that functional circadian rhythm is required for IF-mediated metabolic impact. Our study provides new insights into the role of APC circadian rhythm in regulating WAT fibrosis and metabolic health.

## RESULTS

### Age and chronic high-fat diet increases pro-fibrotic adipose progenitor cells and dampens circadian rhythm in the visceral adipose tissue

To establish an age- and diet-associated adipose dysfunction model, we fed 8-week-old male mice with a 45% high fat diet (HFD) for 10 months prior to the IF regimen. By the end of the HFD regimen, these animals reached 12 months of age, at which point both age and chronic HFD contribute to their metabolic dysfunction (Fig. 1A). As expected, the aged and diet-induced obese (Aged DIO) animals exhibited severe metabolic phenotypes compared to lean young mice (Young), showing higher body weight and impaired glucose clearance (Fig. 1B, 1C). The aged DIO mice showed significantly exacerbated metabolic abnormalities and liver steatosis, compared to the mice fed with 45% HFD for 4 months (DIO) and aged mice fed a normal diet (ND) (Aged) (Fig. S1A-D). Importantly, the WAT of the aged DIO animals exhibited enlarged adipocytes and pro-inflammatory crown-like structures (yellow arrowheads, Fig. 1D), along with significantly increased expression of fibrosis- and inflammation-related genes (Fig. 1E, 1F) and reduced expression of adipogenic genes in the perigonadal WAT (PWAT) (Fig. 1G).

**Figure 1.**
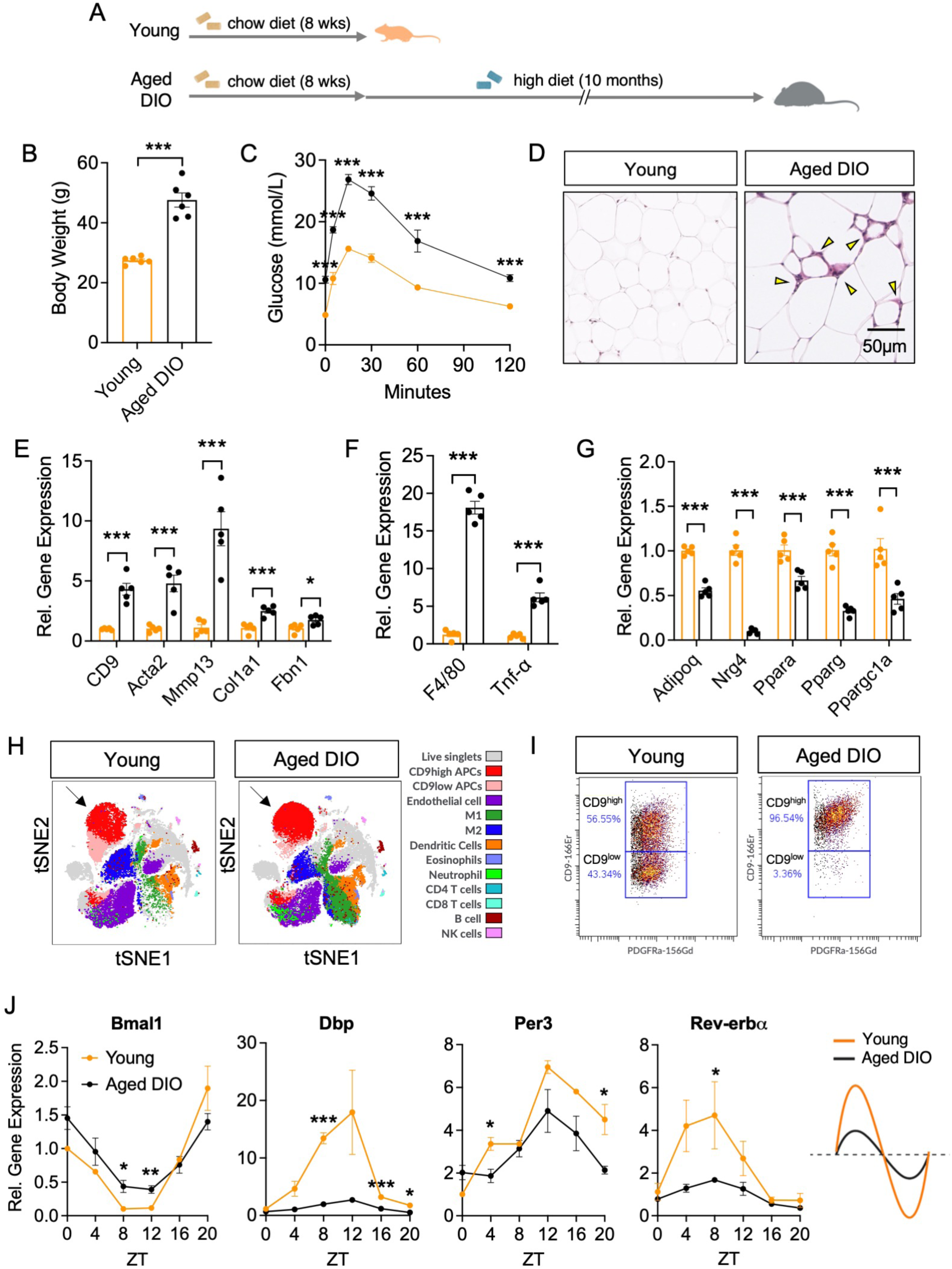
Chronic high-fat diet and ageing results in WAT fibrosis and circadian clock dysfunction. (A) Schematic diagram of aged DIO mouse model. (B) Body weight of young (n = 6) and aged DIO (n=5) mice. (C) IPGTT in young and aged DIO mice. (D) H&E stained images of PWAT from young and aged DIO mice. Yellow arrowheads indicate inflammatory, crown-like structure regions. (E) mRNA levels of fibrotic genes, (F) Inflammatory genes, and (G) Adipogenic genes in PWAT of young and aged DIO mice. (H) Representative tSNE plots of PWAT from young and aged DIO mice illustrating immune and stromal cell composition. Arrows indicate CD9 CD9^high^ APCs. (I) Proportions of CD9^low^ and CD9^high^ APCs of young and aged DIO PWAT. (J) Transcript levels of circadian clock genes in PWAT of young and aged DIO PWAT. Tissues were harvested every 4 hours (n=3 for each ZT). Values are presented as means ± SEM. \**p*<0.05, \*\**p*<0.01, \*\*\**p*<0.005

To gain insight into the cellular changes in WAT by age and chronic HFD, we profiled the composition of the stromal vascular fraction collected from PWAT in young and aged DIO mice, using cytometry by time-of-flight spectrometry (CyTOF) (Fig. 1H) (Lee et al., 2022). In alignment with increased fibrotic gene signatures in WAT (Fig. 1E), the aged DIO PWAT displayed a significantly higher proportion of the pro-fibrotic CD9^high^ PDGFRα+ APCs, with near depletion of the pro-adipogenic CD9^low^ APC subset (Fig. 1I). This suggested that the accumulation of CD9^high^ APCs is the cellular origin of fibrosis in the WAT of aged DIO mice. Aged DIO PWAT also displayed a phenotypic switch in macrophages towards increased pro-inflammatory M1 macrophages and decreased anti-inflammatory M2 macrophages (Fig. S1E). In addition, aged DIO PWAT also contained significantly reduced number of type 2 immune cells, including eosinophils (Fig. S1F) and ILC2 (Fig. S1G), contributing to a pro-inflammatory tissue microenvironment.

Ageing and obesity have been shown to dampen the rhythmicity of core clock genes in adipose tissues (Maury *et al*., 2021; Ribas-Latre *et al*., 2021), and circadian dysregulation is directly linked with the development of fibrosis in lung and multiple tissues (Barbato et al., 2019; Cunningham *et al*., 2020; Rainer, 2020). Thus, we speculated that WAT fibrosis may be associated with circadian misalignment in the WAT. To test this, we harvested PWAT from young and aged DIO mice every 4 hours and assessed the rhythmic expression of clock genes. We analyzed *Bmal1*, *Dbp*, *Per3*, and *Rev-erbα*, which have previously been shown to exhibit robust rhythmicity in WAT. Interestingly, the PWAT of the aged DIO animals displayed significantly dampened amplitudes of the clock genes (Fig. 1J). The rhythmic amplitude of *Bmal1* expression was reduced and *Dbp* and *Rev-erbα* expressions were dampened across all timepoints in the aged DIO PWAT. *Per3* expression was significantly reduced at ZT4 and ZT20 in aged DIO PWAT. Collectively, these data suggest that aged DIO mice represent a model of advanced and severe metabolic phenotypes, along with WAT fibrosis and circadian dampening that can serve as a suitable cohort to test the therapeutic efficacy of IF.

### Intermittent fasting improves age- and diet-induced obesity and diabetic phenotypes

To test the therapeutic potential of IF, the aged DIO animals were subjected to 6 weeks of 2:1 IF (1 day of fasting followed by 2 days of *ad libitum [AL]* feeding) with comparable accumulative food intake (Figs. 2A, S2A) (Kim *et al*., 2017; Kim et al., 2019). After 6 weeks of IF, aged DIO-IF animals exhibited lower body weight compared to aged DIO-AL animals (Fig. 2B). Body composition analysis revealed that the IF-mediated reduction in body weight is due to selective reduction in fat mass without significant changes in lean mass (Fig. 2C). In alignment with this, both inguinal WAT (IWAT) and PWAT mass were significantly reduced after IF (Fig. 2D). Brown adipose tissue (BAT) mass was comparable between the two groups, but liver mass was significantly reduced by IF (Fig. S2B). Histological analysis revealed that the adipocyte size was significantly reduced in both WAT depots (Fig. 2E, 2F) and lipid accumulation was significantly reduced in the liver of the aged DIO-IF mice (Fig. S2C). A key metabolic benefit conferred by IF is an improvement in glucose and insulin homeostasis (Kim *et al*., 2017). As aged DIO animals exhibited hyperglycemia and severe insulin resistance, we assessed whether IF could improve these diabetic phenotypes. While aged DIO-AL and aged DIO-IF animals exhibited a similar levels of glucose intolerance and insulin resistance prior to IF (Fig. 2G, 2I), the mice subjected to 6 weeks of IF displayed significant improvement in both (Fig. 2H, 2J), along with lower fasting insulin levels and lower HOMA-IR (homeostatic assessment of insulin resistance) (Fig. S2D, S2E).

**Figure 2.**
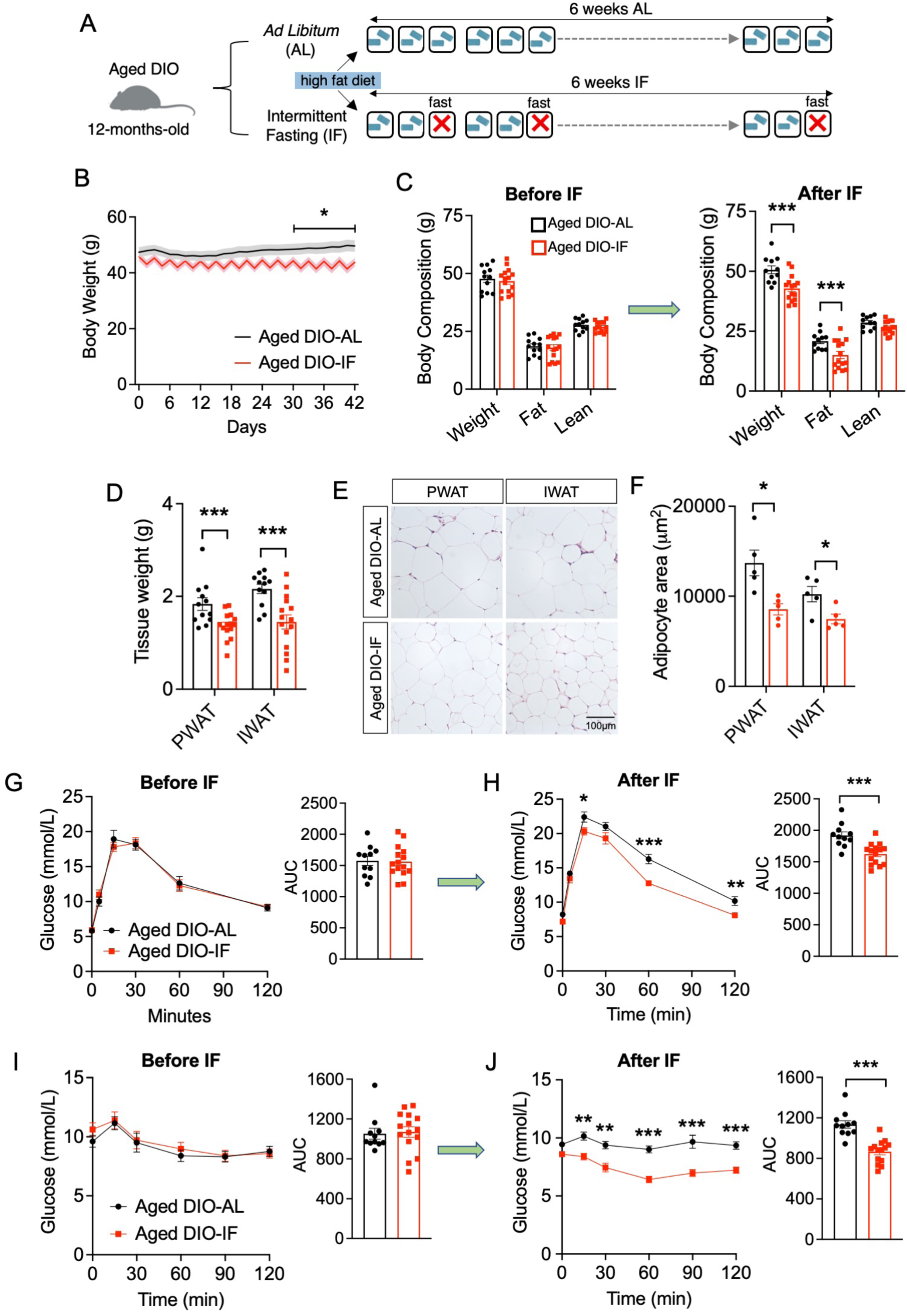
Intermittent fasting reduces fat mass and improves glucose/insulin homeostasis in aged DIO mice. (A) Schematic diagram of IF in aged DIO mice. (B) Body weight of aged DIO-AL (n=12) and aged DIO-IF (n=15) mice during 6 weeks of IF regimen. (C) Body composition analysis before and after 6 weeks of IF. (D) WAT weights after 6 weeks of IF. (E) H&E images of PWAT and IWAT of aged DIO-AL and aged DIO-IF mice. (F) Average adipocyte size of PWAT and IWAT. (G) IPGTT before IF. (H) IPGTT after IF. (I) IPITT before IF. (J) IPITT after IF. Values are presented as means ± SEM. \**p*<0.05, \*\**p*<0.01, \*\*\**p*<0.005

### Intermittent fasting does not enhance adipose thermogenesis in aged DIO animals

Several studies have reported that dietary regimens, such as calorie restriction (CR) and IF, lead to increased non-shivering thermogenesis or browning of WAT, resulting in heightened energy expenditure (Kim *et al*., 2017; Li *et al*., 2017; Martinez-Lopez *et al*., 2017). In particular, IF-subjected mice displayed enhanced diet-induced thermogenesis, where they exhibited increased energy expenditure during the feeding days. Therefore, to test whether WAT browning was the underlying mechanism of IF-mediated metabolic benefits in aged DIO mice, we measured energy expenditure via indirect calorimetric analysis. However, in contrast to previous studies in younger animals (Kim *et al*., 2017; Li *et al*., 2017), aged DIO-IF animals did not display an increased oxygen consumption during the feeding period (Fig. S3A, S3B). Aged DIO-IF animals exhibited significantly reduced energy expenditure during the fasting day and comparable energy expenditure during the feeding days. When total energy expenditure was measured during single cycle of IF (1 day of fasting and 2 days of feeding), aged DIO-IF animals displayed a mildly reduced oxygen consumption rate (Fig. S3C). This was consistently observed when the oxygen consumption rate was analyzed with body weight as a covariate (Fig. S3D) (Mina et al., 2018), further supporting the notion that IF-mediated metabolic benefits are not mediated through increased energy expenditure in aged DIO mice. Histologically, although adipocyte sizes were reduced in the WAT depots, we did not observe regions that resembled browning adipocytes in aged DIO-IF WAT (Fig. 2E, 2F). In alignment with this data, thermogenic gene signatures (i.e., *Ucp1*, *Pgc1α)* were not upregulated in PWAT (Fig. S3E) or brown adipose tissue (BAT) (Fig. S3F) of aged DIO-IF mice, although *Adrb3* expression was significantly increased, which is indicative of activated sympathetic nervous system by fasting (Fig. S3E). These data further validated that 6 weeks of IF may not be sufficient to promote thermogenic activation in aged DIO animals.

### Intermittent fasting resolves adipose tissue fibrosis and enhances plasticity of adipose progenitor cells in aged DIO mice

Aged DIO mice subjected to IF exhibited improved metabolic parameters in the absence of enhanced adipose tissue thermogenesis, suggesting that adaptive thermogenesis is not the main mechanism by which IF confers metabolic benefits in aged DIO animals. Thus, we investigated other mechanisms that could underly improved metabolic function in these mice. A hallmark of adipose tissue dysfunction, WAT fibrosis is closely associated with insulin resistance and impaired metabolic function (Divoux et al., 2010; Halberg et al., 2009; Henegar et al., 2008). CD9^high^ APCs express high levels of collagens and cross-linking enzymes but low levels of genes associated with adipogenesis with fibrogenic phenotype (Hepler *et al*., 2018; Marcelin *et al*., 2017). Similarly, the WAT of aged DIO mice exhibited drastically higher fibrotic gene signatures (Fig. 1E), along with a significantly higher proportion of pro-fibrotic CD9^high^ APCs compared to the lean mice (Fig. 1H, 1I). In turn, the depletion of pro-adipogenic CD9^low^ APCs may contribute to impaired adipogenic and thermogenic function (Hepler *et al*., 2018; Marcelin *et al*., 2017). Hence, we assessed whether IF-mediated metabolic benefits may be mediated through the resolution of WAT fibrosis. Compared to AL mice, the PWAT of aged DIO-IF mice had reduced fibrosis (Picrosirius staining) (Fig. 3A) and displayed significant downregulation of fibrotic markers, including *Col1a1*, *Acta2*, *Mmp13* and *CD9* (Fig. 3B). Additionally, we observed a significant reduction in senescence-associated markers, such as *p21* and *p53* (Fig. 3B), which have been associated with reduced cellular plasticity and impaired beige adipocyte formation (Berry *et al*., 2017). To investigate whether WAT fibrosis is associated with the phenotypic alteration of APCs, we performed single cell mass cytometry (CyTOF) and an APC differentiation assay (Fig. 3C). In alignment with gene expression data, the CyTOF analysis of stromal cells revealed significantly reduced CD9^high^ (pro-fibrotic) and increased CD9^low^ (pro-adipogenic) APCs in the PWAT of aged DIO-IF animals (Fig. 3D). To further investigate whether IF improves the adipogenic potential of APCs, which has been shown to be reduced in aged WAT (Berry *et al*., 2017), we isolated PDGFRα+ APCs from aged DIO-AL and aged DIO-IF PWAT and differentiated them *in vitro* (Fig. 3E). At day 8 of adipogenic differentiation, we observed a significantly higher lipid droplet accumulation in aged DIO-IF mice (Fig. 3E, 3F), suggesting that APCs from aged DIO-IF mice are more prone to healthy adipose remodeling via *de novo* adipogenesis (Pellegrinelli *et al*., 2016; Rutkowski *et al*., 2015). This finding was in line with the characteristics of IF-treated WAT, with significant upregulation of adipogenic and healthy adipokine gene signatures, including *Nrg4*, *Pparg*, and *Ppara* (Fig. 3G). In particular, *Nrg4* has been reported to attenuate HFD-induced liver steatosis (Wang et al., 2014), suggesting that the production of healthy adipokines in aged DIO-IF mice may alleviate the fatty liver phenotype induced by age and chronic HFD. Collectively, these data suggested that IF-induced reprogramming of APCs promotes healthy adipogenesis and suppresses WAT fibrosis. These data implies that even in the absence of WAT thermogenesis, the anti-fibrotic effects of IF may confer metabolic benefits to aged DIO mice.

**Figure 3.**
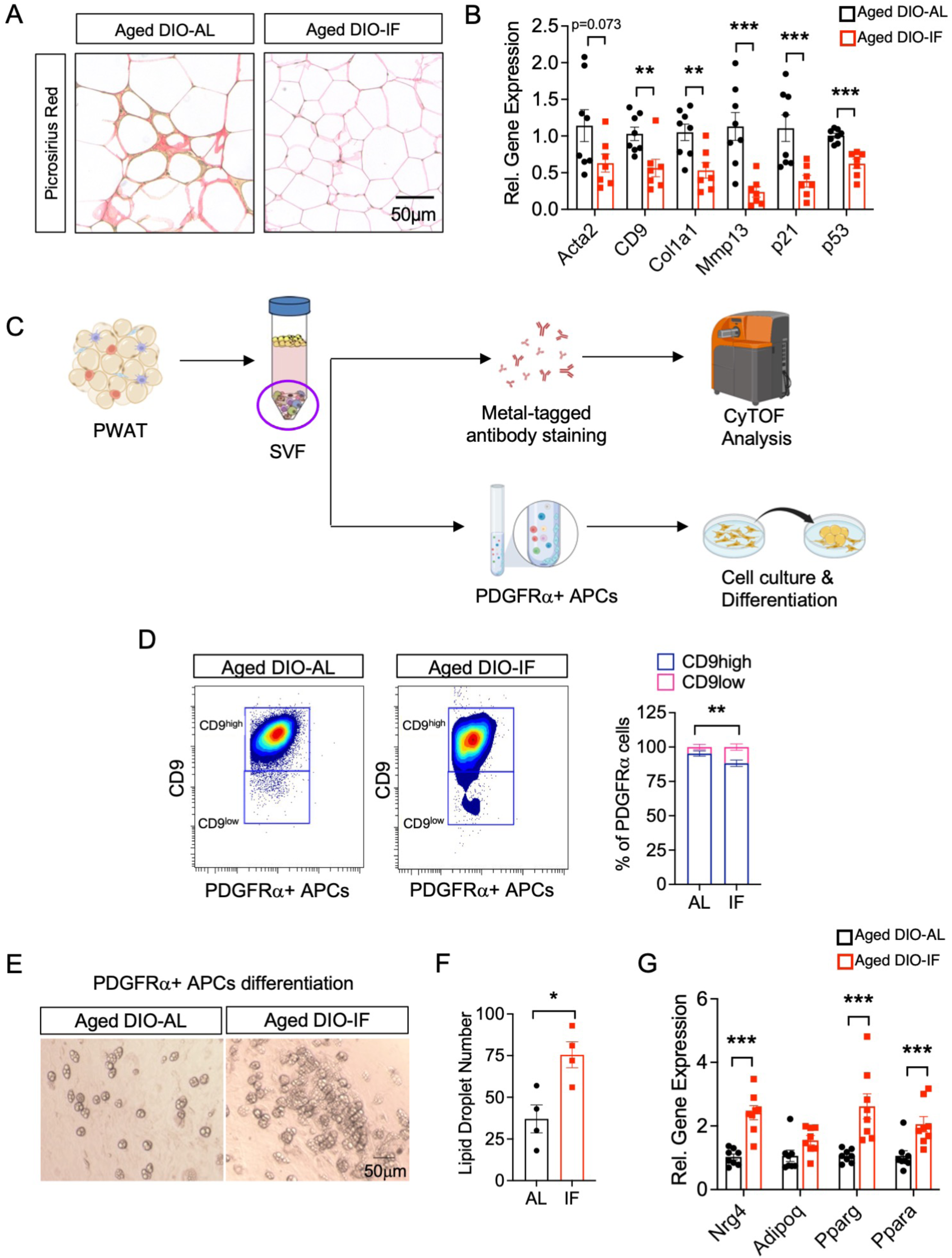
Intermittent fasting reduces adipose tissue fibrosis. (A) Picrosirius red staining of PWAT from aged DIO-AL and aged DIO-IF mice. (B) mRNA levels of fibrosis genes in PWAT. (C) Schematic diagram of assessing APC phenotype using CyTOF analysis and *in vitro* cell culture and differentiation. (D) Characterization of CD9^low^ and CD9^high^ PDGFRα+ APCs in PWAT. (E) *In vitro* adipogenic differentiation of APCs from aged DIO-AL and aged DIO-IF PWAT. Images were taken at day 8 post-differentiation. (F) Number of lipid droplets in differentiated APCs. (G) mRNA levels of adipogenic gene signatures in PWAT. Values are presented as means ± SEM. \**p*<0.05, \*\**p*<0.01, \*\*\**p*<0.005

### Single nuclei RNA sequencing reveals IF-mediated circadian changes in the APCs

Although our gene expression and CyTOF data suggested that IF may confer metabolic benefits through anti-fibrotic remodeling of APCs, the underlying mechanisms of how IF regulates APCs are unknown. Thus, to gain insight into the cellular targets of IF and underlying pathways that are altered by IF, we performed single nuclei RNA sequencing (snRNA-seq), which allows analysis of both stromal cell populations and mature adipocytes. Unsupervised clustering revealed 14 distinct cell types including: mature adipocytes (marked by high expression of *Plin1* and *Dgat2*), macrophages (*Itgam*, *Adgre1*, and *Mrc1*), endothelial cells (*Pecam1* and *Flt1*), lymphocytes (*CD79a*, *CD226*, and *CD247*), mesothelium (Bnc1 and *Upk3b*), and APCs (*PDGFR*α and *CD34*) (Figs. 4A, S4A). A common mesenchymal progenitor cell marker, *PDGFRα,* was expressed in both APC clusters (APC1 and APC2), and other APC markers, including *PDGFRβ, CD34, CD29, DPP4*, and *Sca-1* (*Ly6a*), and *CD9*, were expressed in one or both of APC subsets (Fig. S4B). Proportional cellular frequencies (normalized by total number of specific cell populations) showed major differences in the cell numbers of adipocytes, APCs, and macrophages (Fig. 4B). Cell type-specific transcriptomic changes in response to IF were assessed using differential gene expression and gene ontology (GO) enrichment analysis. To visualize comprehensive molecular changes across different cellular clusters, we grouped networks based on enriched GO terms that were functionally related (Fig. 4C, 4D). Biological processes that were upregulated in aged DIO-IF included rhythmic process, fat cell differentiation, and synapse assembly (Fig. 4C). Notably, circadian rhythm related biological processes were upregulated in APCs, mesothelial cells, and macrophages. Downregulated GO terms in aged DIO-IF included extracellular matrix (ECM) organization, lymphocyte differentiation and activation of leukocyte chemotaxis, and antigen processing and presentation (Fig. 4D). Pathways associated with ECM organization were specifically downregulated in APC clusters, while suppression of immune response and chemotaxis was downregulated across APC, macrophages, mesothelial cells, and B cells. The APC2 cluster presented with a strong dichotomy of increased expression of circadian rhythm-related genes and decreased expression of ECM organization and collagen-related processes (Fig. 4E). Correspondingly, the expression of various ECM-associated genes, such as *Col1a1*, *Col6a1*, *Fbn1*, and *Fn1*, was significantly reduced within the APC clusters (Fig. 4F). Intriguingly, mature adipocytes of aged DIO-IF PWAT had upregulated fatty acid metabolism and thermogenesis-associated pathways, while cell growth and cell junction assembly pathways were downregulated (Fig. 4G). This result is consistent with the recent study that illustrated increased diet-induced thermogenesis in mature adipocytes by time-restricted feeding (Hepler et al., 2022). In addition, genes associated with healthy adipokines were also increased in aged DIO-IF adipocytes (Fig. 4H). Macrophages in aged DIO-AL were enriched in various inflammatory pathways, such as cell chemotaxis, antigen processing and presentation, and chemokine-mediated signaling pathways, while macrophages in aged DIO-IF were enriched for circadian rhythm and hormone-mediated signaling pathways (Fig. S4C, S4D). Furthermore, using gene activity scores derived from genes belonging to specific biological processes, we confirmed that circadian rhythm-related genes were significantly upregulated across multiple cell types (Fig. 4I), including macrophages/monocytes, mesothelium, adipocytes, B cells, and APCs, with APCs displaying the most significant changes (Fig. 4J). In contrast, the gene activity scores for collagen biosynthesis were significantly downregulated in the mesothelial cells and APCs (Fig. 4K) with the most significant changes observed in APC (Fig. 4L). This dichotomy strongly suggested that the anti-fibrotic effects of APCs were accompanied by the upregulation of circadian rhythm pathways. Next, we identified influential transcription factors with gene targets that were significantly altered by IF, based on literature aggregated ChIP-seq databases (Kuleshov et al., 2016; Lachmann et al., 2010) (Fig. 4M). We identified transcription factors that are critical circadian regulators, including CLOCK and BMAL1, which were statistically significant within the APC clusters. Since circadian rhythm was revealed as one of the most significantly altered pathways in WAT by IF, we harvested PWAT from the aged DIO-AL and aged DIO-IF mice every 4 hours to assess how the rhythmicity of clock genes is altered by IF (Fig. 4N). Importantly, we observed a significant increase in the amplitudes and mesor (rhythm-adjusted mean) of *Dbp* and *Per3* expression by IF treatment, with significantly higher peaks at ZT8 and ZT12, respectively. Additionally, we observed a significant increase in the amplitude of *Bmal1* expression, with a significantly lower trough at ZT8. While the amplitude of *Rev-erbα* was unchanged by IF, a phase shift was observed, which may have been influenced by slight differences in feeding times, as IF mice consumed food during not only the active period but also the inactive phase after fasting day.

**Figure 4.**
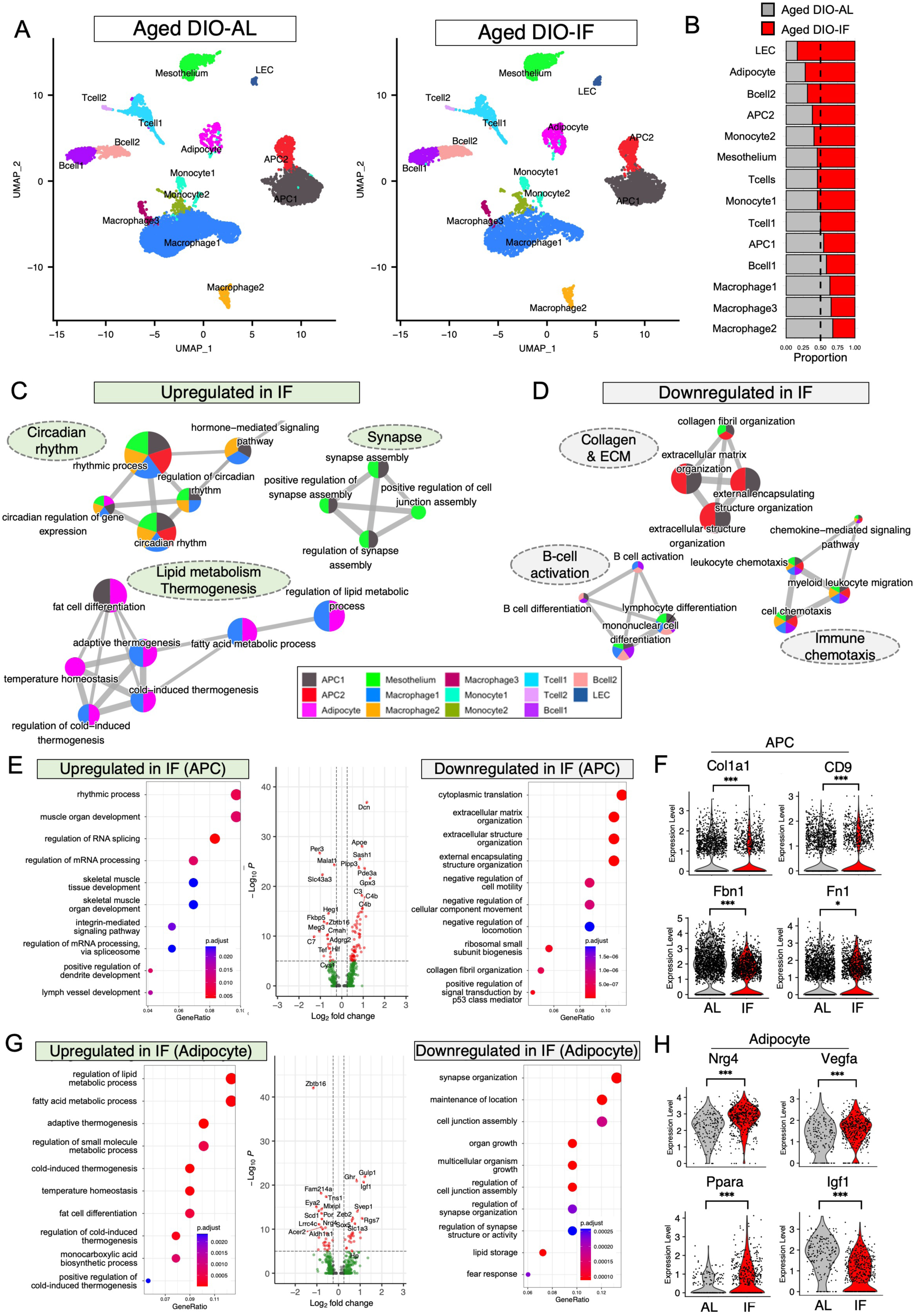
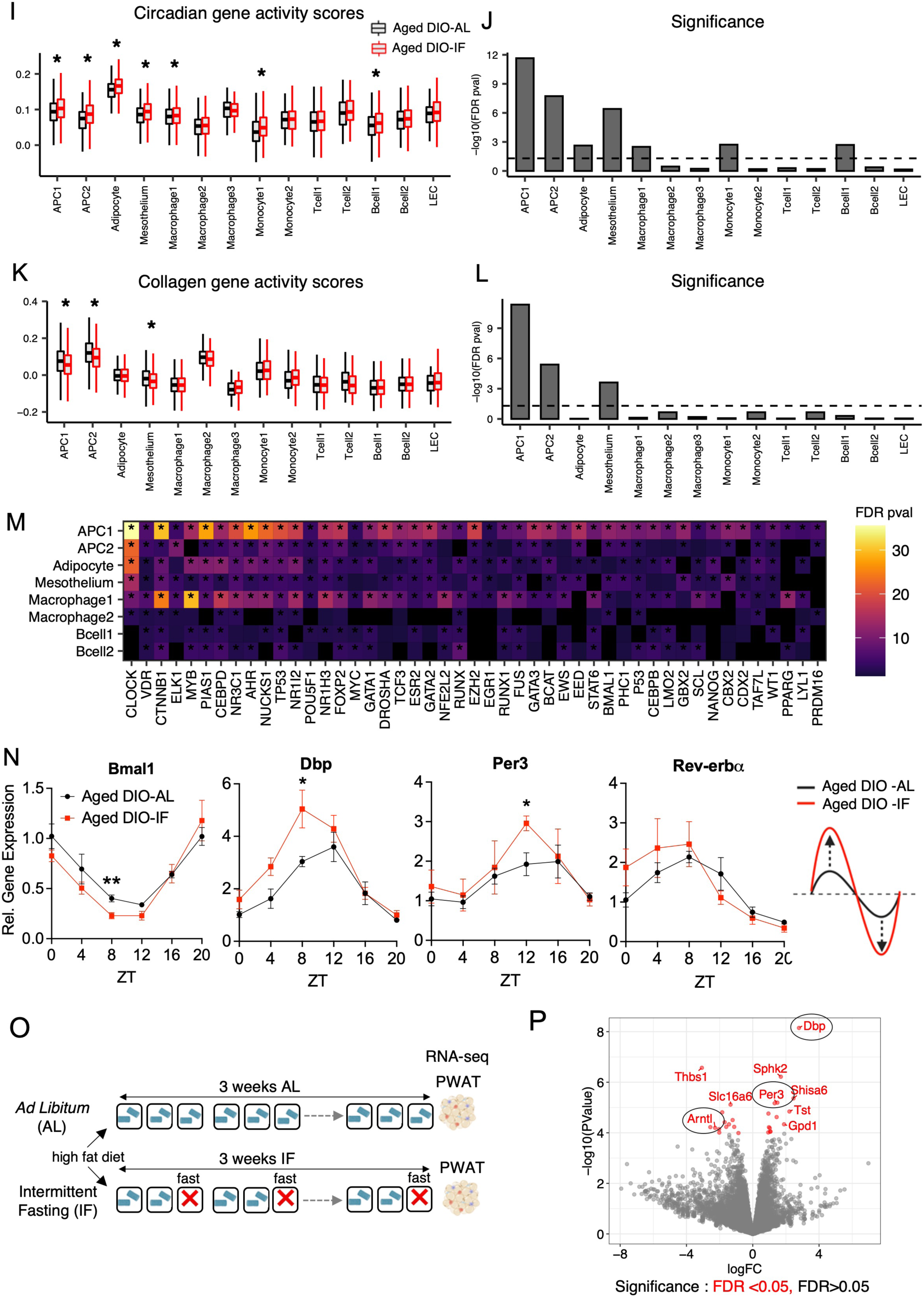
Single nuclei sequencing reveals alteration of circadian pathways in WAT by IF. (A) Representative UMAP plots of PWAT harvested from aged DIO-AL and aged DIO-IF mice. PWAT of 3 mice were pooled for each sample. (B) Proportional frequencies of cell types in PWAT, normalized by total number of cells. (C) Gene ontology (GO) networks of significantly upregulated DEGs in aged DIO-IF PWAT. (D) GO networks of significantly downregulated DEGs in aged DIO-IF PWAT. (E) Significantly altered GO terms in APC cluster. (F) Transcript levels of fibrotic markers in APCs. (G) Significantly altered GO term pathway in adipocytes. (H) Transcript levels of adipogenic markers in adipocytes. (I) Module scoring of circadian gene sets across cell clusters. (J) Statistical significance of module scoring of circadian genes. (K) Module soring of collagen biosynthesis gene sets across cell clusters. (L) Statistical significance of module scoring of collagen biosynthesis genes. (M) Transcription factors that have been identified by published ChIP-seq datasets, that modulate significantly altered genes by IF. (N) Expression levels of circadian genes in PWAT harvested every 4 hours (n = 3-4 per ZT). (O) Schematic diagram of short-term IF and tissue harvesting timepoint. (P) Volcano plot displaying top differentially regulated genes by short-term IF. Genes in red indicate significance of FDR <0.05. Values are presented as means ± SEM. \**p*<0.05, \*\**p*<0.01, \*\*\**p*<0.005

Despite significant alterations in the circadian pathway by IF, these could be merely a consequence of body or fat weight loss and metabolic improvement by IF, not a driver of metabolic benefits. Thus, to test this, we harvested PWAT from mice that were subjected to 3 weeks of IF or AL and performed bulk-tissue RNA sequencing (Fig. 4O). Here, the animals did not display any metabolic improvements; no difference was observed in body weight and composition, tissue weight, or fasting glucose (Fig. S4E-H). This allowed us to identify pathways that are altered earlier on during the IF regimen. Among the top differentially regulated pathways, we identified upregulation of mitochondrial activity, fatty acid metabolism, thermogenesis, and circadian pathways (Fig. S4I). Moreover, we identified key circadian genes (*Bmal1*, *Dbp*, *Per3*) as some of the top differentially expressed genes in short-term IF PWAT (Figs. 4P and S4J), further supporting that impact on circadian pathways may be early drivers of IF-mediated metabolic improvements.

### Animals with systemic and APC-specific *Bmal1* deletions demonstrate WAT fibrosis

The association between circadian dysfunction and metabolic impairment has been well documented. *Bmal1* knockout in adipocytes leads to increased weight gain, adipocyte hypertrophy, dampened energy expenditure, and altered fatty acid metabolism (Paschos *et al*., 2012). Our data identified the circadian pathway as a potential mediator of IF-induced metabolic benefits through suppression of WAT fibrosis. Conversely, circadian dysfunction may drive WAT fibrosis and metabolic abnormalities. However, whether circadian rhythm or the core clock gene *Bmal1* is associated with fibrotic activation in WAT is unknown. To test this, we assessed the WAT phenotype of whole-body *Bmal1* knockout (*Bmal1* KO) mice (Fig. 5A). Consistent with previous studies (Kondratov *et al*., 2006; Laermans et al., 2015), *Bmal1* KO mice completely lost the diurnal rhythm of energy expenditure with mildly reduced total oxygen consumption (Fig. S5A–C). Although *Bma1* KO animals did not display a significant difference in body weight (Fig. 5B), fat composition and adiposity were greater than control mice, with increased adipocyte size in both the PWAT and IWAT of *Bmal1* KO mice (Figs. 5C-F and S5D). Importantly, the WAT from *Bmal1* KO animals exhibited increased fibrotic signatures (Fig. 5G), even with normal chow diet feeding, which was in alignment with the increased CD9 median intensity within PDGFRα+ APCs (Fig. S5E). This supports that circadian disruption can cause fibrotic progression in WAT. To gain insight into the cell populations that contribute to increased fibrotic signatures, we performed snRNA-seq for WAT of *Bmal1* control (Ctrl) and KO mice. Similar to snRNA-seq on the aged DIO WAT, unsupervised clustering identified 12 distinct cell types (Fig. 5H). Interestingly, we found that both APC clusters were proportionally reduced in *Bmal1* KO WAT compared to control WAT (Fig. 5I). Importantly, APC clusters exhibited increased signatures of extracellular matrix organization and fibrosis-related signatures, with increased *Col1a1* and *Col6a1* expressions (Fig. 5J, 5K). In contrast, decreased signatures of healthy adipokine and thermogenesis were observed in mature adipocyte clusters of *Bmal1* KO mice (Fig. 5L, 5M), further supporting that *Bmal1* KO mice display unhealthy WAT phenotypes with increased fibrosis signatures.

**Figure 5.**
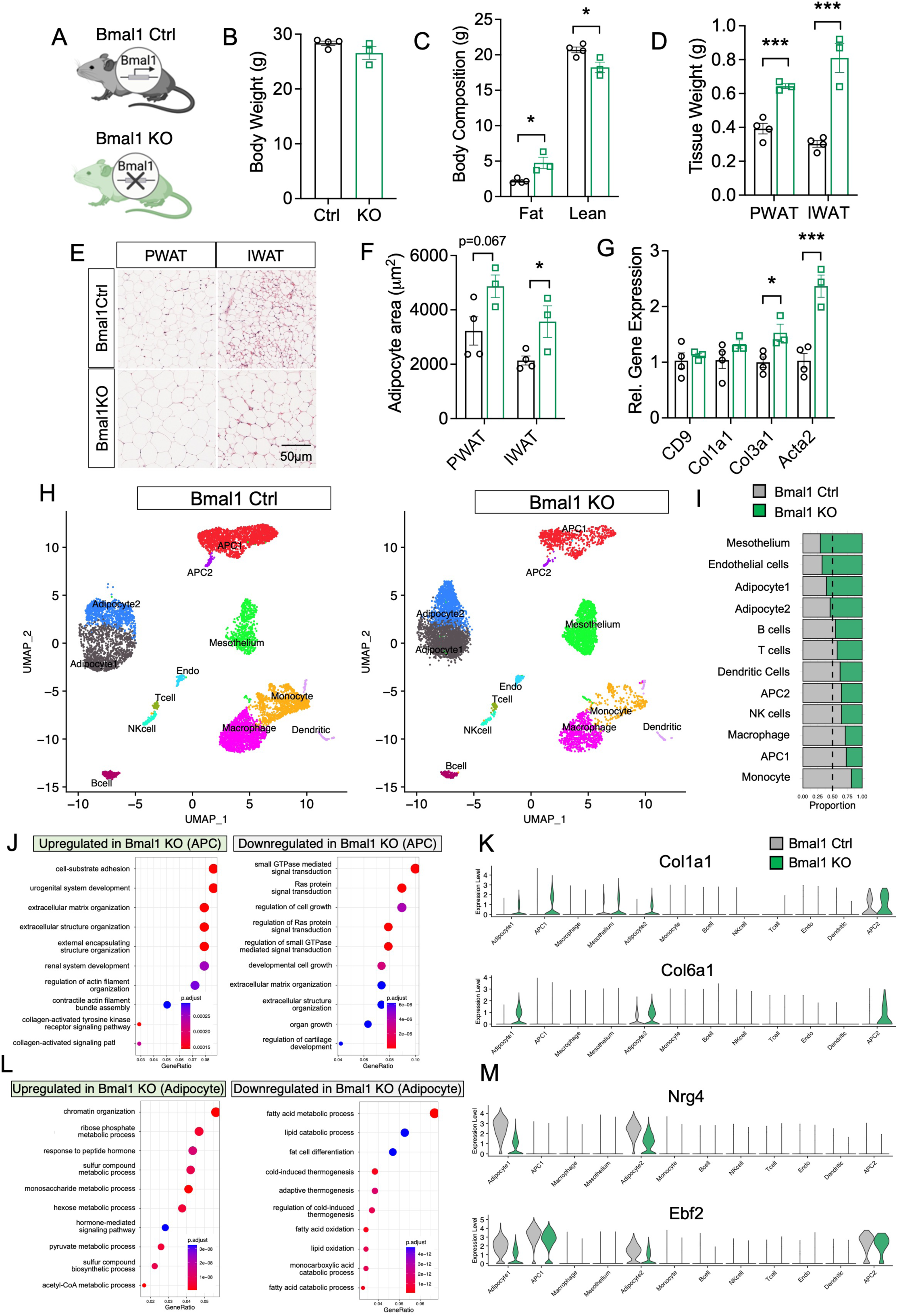
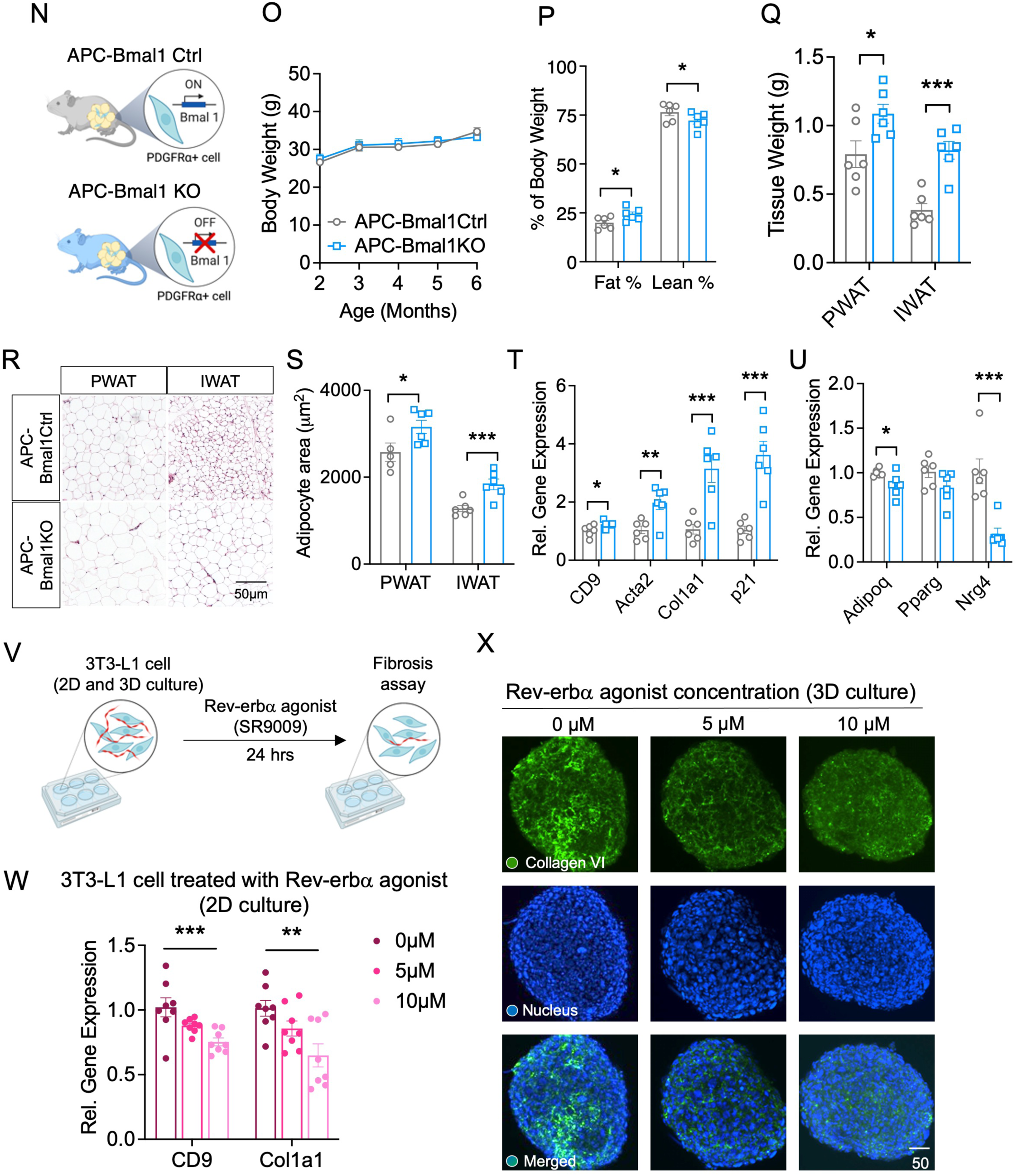
Mice with global and APC-specific *Bmal1* deficiency exhibit increased fat mass and adipose tissue fibrosis. (A) Schematic diagram of *Bmal1* Ctrl and whole-body *Bmal1* KO mouse model. (B) Body weight of *Bmal1* Ctrl (n = 4) and *Bmal1* KO (n = 3) mice at 8 weeks of age. (C) Body composition showing fat and lean mass. (D) WAT weights. (E) H&E stained images of WAT. (E) Adipocyte size. (G) mRNA levels of fibrosis genes in PWAT. (H) Representative UMAP plots of PWAT from *Bmal1* Ctrl and *Bmal1* KO mice. (I) Proportional frequencies of cell types in PWAT, normalized by the total number of cells. (J) Significantly altered GO terms in APC cluster. (K) Violin plots displaying *Col1a1* and *Col6a1* transcript levels across cell clusters. (L) Significantly altered GO terms in adipocytes. (M) Violin plots displaying *Nrg4* and *Ebf2* transcript levels across cell clusters. (N) Schematic diagram of APC-*Bmal1* Ctrl and APC-*Bmal1* KO mouse model. (O) Body weight on ND from 2 to 6 months of age. (P) Percentage of fat and lean mass, normalized by body weight. (Q) WAT weights. (R) H&E stained images of WAT. (S) Adipocyte size. (T) mRNA levels of fibrosis genes in PWAT. (U) mRNA levels of adipogenic genes in PWAT. (V) Schematic diagram of *in vitro* SR9009 treatment in 3T3-L1 cells. (W) mRNA levels of *CD9* and *Col1a1* in cells treated with SR9009. (X) Immunofluorescence staining of collagen type VI in 3D 3T3-L1 spheroids. Values are presented as means ± SEM. \**p*<0.05, \*\**p*<0.01, \*\*\**p*<0.005

Although the *Bmal1* KO adipose tissue data support our hypothesis that loss of circadian rhythm may drive WAT fibrosis, these results may be a consequence of the loss of behavioural rhythmicity due to a globally dysfunctional circadian clock in *Bmal1* KO mice. Thus, to address the role of the circadian clock of APC in the pathogenesis of WAT fibrosis, we generated APC-specific *Bmal1* KO mice by crossing PDGFRα-Cre mice with *Bmal1* floxed mice (PDGFRα-Cre:*Bmal1*^flox/flox^, APC-*Bmal1* KO) (Fig. 5N). Littermate mice homozygous for *Bmal1* allele but not carrying Cre recombinase were used as control mice (APC-*Bmal1* Ctrl). We found a significant reduction in the expression of *Bmal1 and Dbp,* a target gene of *Bmal1*, within APCs and adipocytes of APC-*Bmal1* KO mice, but not in the PDGFRα negative (PDGFRα-) cells, indicating successful knocking out of *Bmal1* within APCs (Fig. S5F, S5G). Unlike *Bmal1* KO mice, APC-*Bmal1* KO mice displayed a partially recovered diurnal rhythm of energy expenditure, with overall energy expenditure comparable to control mice (Fig. S5H-J). APC-*Bmal*1 KO mice did not have significant differences in body weight (Fig. 5O). However, similar to whole-body *Bmal1* KO mice, APC-*Bmal1* KO mice showed a significant increase in proportional fat mass, along with increased size of adipocytes, compared to APC-*Bmal1* Ctrl mice (Fig. 5P-S). Importantly, fibrosis and senescence-associated gene expression was significantly increased in the PWAT of APC-*Bmal1* KO mice (Fig. 5T). In contrast, adipogenic genes, such as *Adipoq* and *Nrg4* (associated with improvement in fatty liver disease), were significantly downregulated (Fig. 5U). Next, to test whether a functional circadian clock is directly associated with anti-fibrotic changes in APCs, we treated murine preadipocytes (3T3-L1) with SR9009, a *Rev-erbα* agonist known to activate circadian rhythms (Dierickx et al., 2019; Sulli et al., 2018), for 24 hours. We then examined the cellular fibrosis signatures and immunostained for collagen type VI. We found that treatment with SR9009 significantly reduced the expression of fibrosis-related signatures (Fig. 5W) and displayed reduced collagen type VI levels (Fig. 5X), suggesting that pharmacological activation of the circadian clock suppresses fibrotic progression in preadipocytes.

### Animals with whole-body or APC-specific *Bmal1* deficiency exhibit a blunted response to IF

Our data revealed that IF induces circadian reprogramming of APCs, which may be associated with their anti-fibrotic effects and metabolic improvement. Interestingly, whole-body and APC-specific *Bmal1* knockout mice exhibited WAT fibrosis, suggesting that circadian disruption promotes unhealthy WAT remodeling and its fibrotic progression. As IF improved glucose homeostasis and WAT remodeling in aged DIO mice via circadian regulation of APCs, we next tested whether a functional clock is required for the metabolic benefits of IF. To test this, we performed 6 weeks of IF in whole body *Bmal1* KO and APC-*Bmal1* KO mice (Fig. 6A). Both whole-body and APC-specific *Bmal1* KO mice showed similar body weight cycling compared to littermate controls during 6 weeks of IF, without significant differences in body weight (Fig. 6B, 6E). However, we found that the adipocyte sizes were significantly reduced in *Bmal1* Ctrl-IF and APC-*Bmal1* Ctrl-IF mice, whereas the degree of size reduction was moderate in the WAT of *Bmal1* KO-IF and APC-*Bmal1* KO-IF (Figs. 6C, 6D, 6F, 6G, S6A, and S6F), demonstrating blunted response of adipocytes to IF in these circadian KO models. To test whether IF-mediated metabolic effects are also affected by *Bmal1* KO, we next assessed insulin sensitivity. While *Bmal1* Ctrl animals showed significant improvement in insulin sensitivity, with a sustained decrease in blood glucose level (Fig. 6H), *Bmal1* KO animals subjected to IF did not demonstrate any improvement in insulin sensitivity (Fig. 6I). This was without any difference in their energy expenditure (Fig. S6B-E). The diminished metabolic effect from IF in *Bmal1* KO mice may be primarily due to their global lack of functional circadian rhythm. To address whether the blunted metabolic response to IF is primarily attributed to lack of APC circadian rhythm, 12-week-old APC-*Bmal1* Ctrl and APC-*Bmal1* KO mice were subjected to 6 weeks of IF. Similar to whole-body *Bmal1* Ctrl and KO mice, while APC-*Bmal1* Ctrl mice subjected to IF displayed significant improvement in insulin tolerance test (Fig. 6J), IF-subjected APC-*Bmal1* KO mice exhibited only a mild improvement in insulin tolerance test (Fig. 6K). This phenotype was again not associated with their energy expenditure changes (Fig. S6G-J). Together, these data suggest a blunted metabolic benefit of IF in mice lacking functional circadian clock in APCs. To investigate whether the blunted insulin sensitivity effect of IF in circadian KOs is associated with lack of resolution of WAT fibrosis, we assessed fibrotic signatures in both whole-body and APC targeted KO models. Notably, expressions of fibrosis-related genes were significantly reduced in the PWAT of *Bmal1* Ctrl mice, however, this effect was diminished in the PWAT of *Bmal1* KO models (Fig. 6L). As APCs are the major source of WAT fibrosis (Marcelin *et al*., 2017), to further clarify the role of functional clock gene in anti-fibrosis effect of IF in a cell-specific manner, APCs were isolated from PWAT of APC-*Bmal1* Ctrl and KO mice. While fibrotic signatures were decreased by IF in APC-*Bmal1* Ctrl group, no significant changes were observed in APC-*Bmal1* KO group (Fig. 6M). This suggests that the anti-fibrotic effects of IF requires functional APC circadian clock, and without a functional clock, IF does not confer an equivalent level of metabolic benefits, such as insulin sensitivity. Collectively, these data suggest that functional circadian clock is an indispensable factor in conferring IF-mediated benefits.

**Figure 6.**
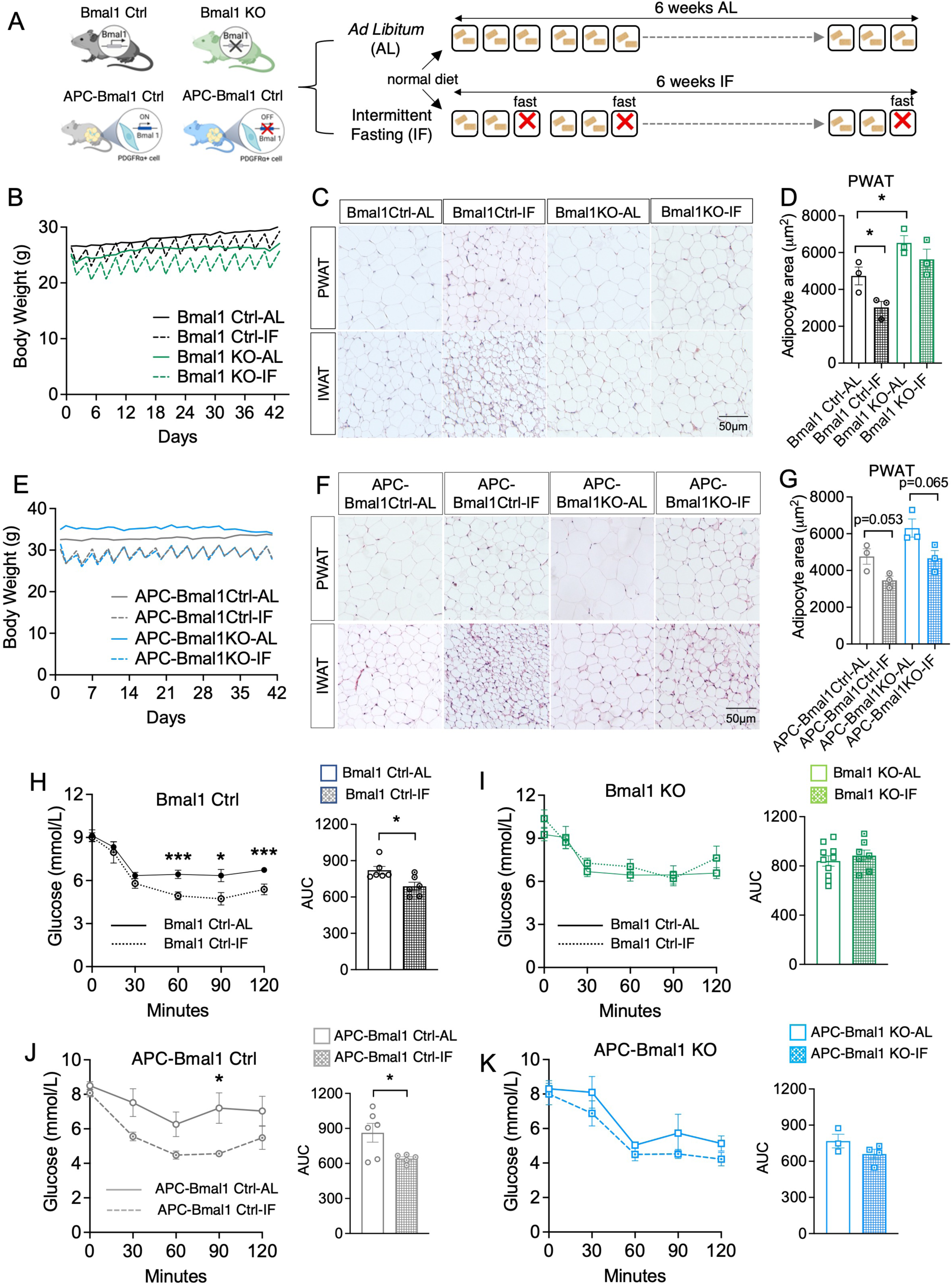
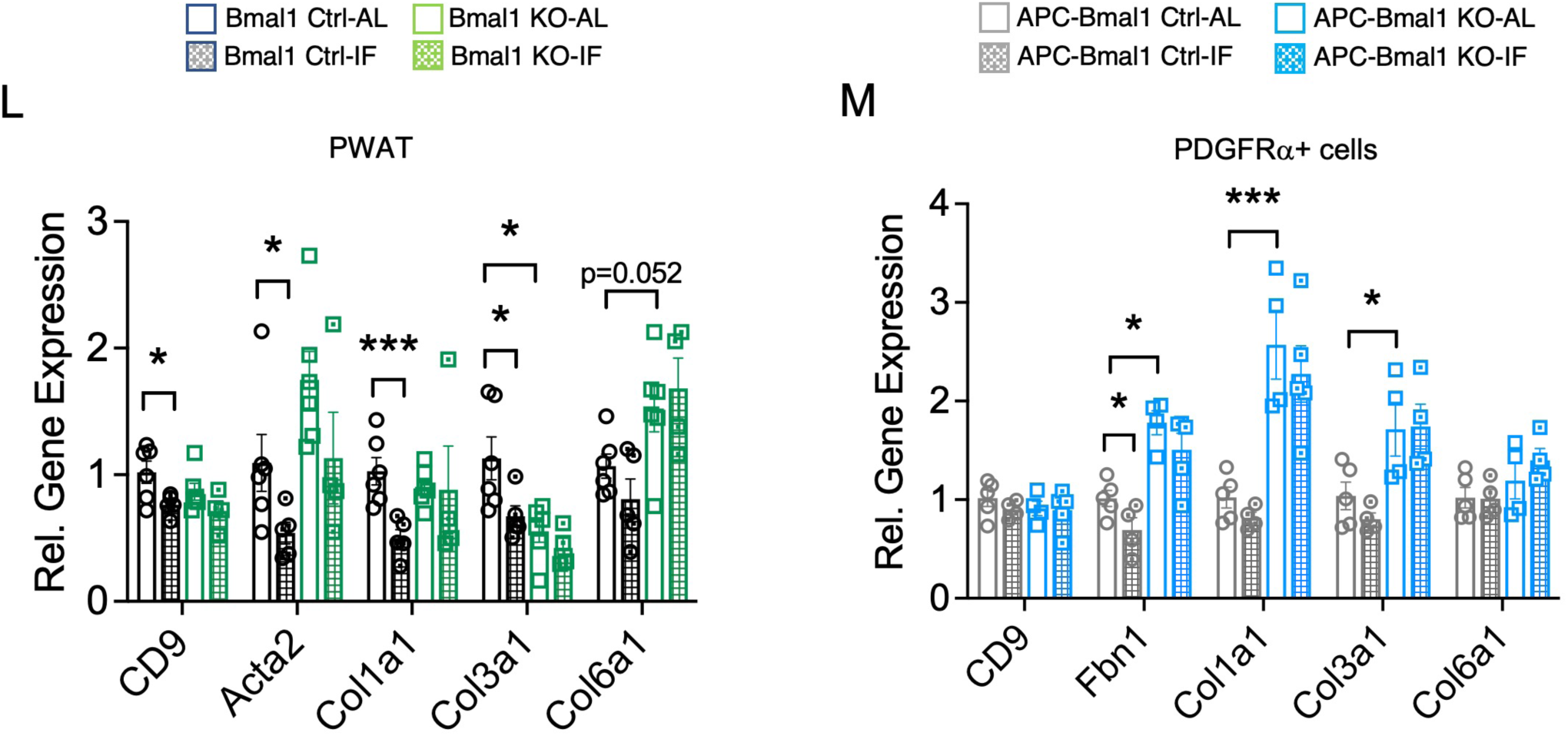
Mice lacking global or APC-specific *Bmal1* exhibit blunted response to intermittent fasting. (A) Schematic diagram of mice subjected to 6 weeks of IF. (B) Body weights of *Bmal1* Ctrl and *Bmal1* KO mice during 6 weeks of IF. (n = 5 for *Bmal1* Ctrl-AL; n = 5 for *Bmal1* Ctrl-IF; n = 6 for *Bmal1* KO-AL; n = 4 for *Bmal1* KO-IF) (C) H&E images of PWAT and IWAT of *Bmal1* Ctrl and *Bmal1* KO mice under AL or IF. (D) Adipocyte size of PWAT (n = 3 for each group). (E) Body weights of APC-*Bmal1* Ctrl and APC-*Bmal1* KO mice during 6 weeks of IF. (n = 6 for APC-*Bmal1* Ctrl-AL, n = 5 for APC-*Bmal1* Ctrl-IF, n = 3 for APC-*Bmal1* KO-AL, and n = 4 for APC-*Bmal1* KO-IF). (F) H&E images of PWAT and IWAT of APC-*Bmal1* Ctrl and APC-*Bmal1* KO mice under AL or IF. (G) Adipocyte size of PWAT (n = 3 for each group). (H) IPITT of *Bmal1* Ctrl mice. (I) IPITT in *Bmal1* KO mice. (J) IPITT of APC-*Bmal1* Ctrl mice. (K) IPITT of APC-*Bmal1* KO mice. (L) mRNA levels of fibrotic genes in PWAT of *Bmal1* Ctrl and *Bmal1* KO mice under AL or IF. (M) mRNA levels of fibrotic genes within PDGFRα+ APCs in PWAT of APC-*Bmal1* Ctrl and APC-*Bmal1* KO mice under AL or IF. \**p*<0.05, \*\**p*<0.01, \*\*\**p*<0.005

### Circadian pathways are altered in APCs in insulin-resistant human visceral adipose tissue

Our *in vivo* and *in vitro* data revealed that a dysfunctional circadian rhythm leads to WAT fibrosis and a blunted response to IF. To assess whether disrupted circadian pathways in WAT are associated with impaired metabolism in humans, we performed snRNA-seq on visceral adipose tissues (VAT) collected from 4 patients undergoing bariatric surgery. Patients were matched for sex, with similar age (Supplementary Table 1). All patients had body mass index (BMI) > 60 kg/m^2^, however two patients were relatively insulin sensitive with lower fasting insulin and HOMA-IR in a healthy range (insulin sensitive group), compared to the other two patients who had evidence of severe insulin resistance (insulin resistant group) (Fig. S7A). We aimed to investigate the cellular and transcriptomic differences in the WAT of these two groups to identify the distinguishing factors that could underly their relative insulin sensitivity, independent of body weight and adiposity. Unsupervised clustering using representative marker genes, resulted in 17 clusters analogous to the clusters found from mouse snRNA-seq in WAT (Fig. 7A). Interestingly, the cellular proportions of the two APC populations, APC1 and APC2, were both higher in insulin-sensitive fat (Fig. 7B). GO term enrichment analysis revealed various biological processes that are significantly different between the two groups. Upregulated processes in the insulin-sensitive WAT included circadian rhythm and response to insulin or corticosteroid across multiple cell clusters, including adipocytes, APCs, macrophages, and mesothelial cells (Fig. 7C). In contrast, upregulation of actin filament organization and cell adhesion was observed in adipocytes, APCs, and macrophages of insulin-resistant WAT (Fig. 7D). In particular, adipocytes of insulin-resistant WAT demonstrated upregulation of the SMAD signaling pathway, known to be associated with increased TGFβ activity and fibrosis. Importantly, cluster-specific GO term enrichment analysis revealed the upregulation of the rhythmic process and response to peptide hormones in insulin sensitive APC populations and adipocytes (Figs. 7E, 7F, and S7B). In contrast, GO term analysis showed the upregulation of cell-matrix adhesion and catabolic processes in insulin resistant APC populations and adipocytes (Figs. 7E, 7F, and S7B). Feature plots revealed that the expression levels of key circadian genes (e.g., *BMAL1*, *CRY1*, *CLOCK*) were significantly dampened across various cell clusters of insulin resistant WAT, including APCs (Figs. 7G). Our mouse data showed a dichotomy of circadian rhythm and fibrosis signature, and pharmacological activation of circadian rhythm using *Rev-erbα* agonist, decreased collagen expression levels in 3D 3T3-L1 spheroids (Figs. 5V-X). Thus, to test whether a functional circadian clock is associated with fibrotic alterations in human APCs, we generated human preadipocyte spheroids (Fig. S7C) (Taylor et al., 2020) and treated them with *Rev-erbα* agonist SR9009. Similar to what we observed in 3D murine preadipocytes, SR9009 treatment reduced the expression of collagen expression (Fig. S7D), suggesting that pharmacological activation of the circadian clock suppresses fibrotic progression in human preadipocytes. These results suggest that the global dampening of circadian rhythm gene expression in human WAT may be associated with WAT fibrosis and adverse metabolic outcomes such as insulin resistance, independent of body weight.

**Figure 7.**
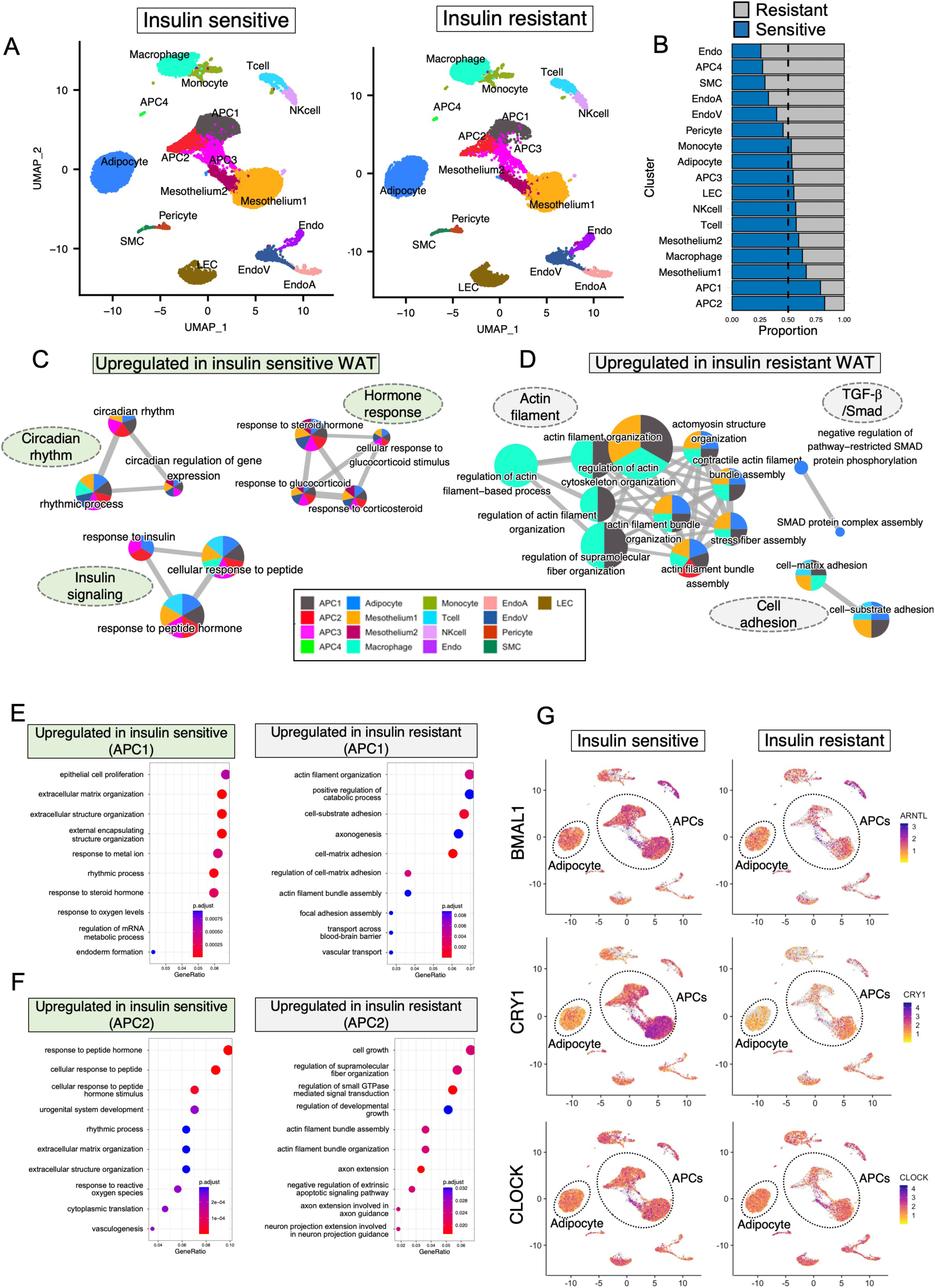
Humans with type 2 diabetes displayed altered circadian pathways in adipocytes and APCs within omental WAT. (A) Representative UMAP plots of omental WAT from insulin-sensitive and insulin-resistant individuals (n = 2 for each group). (B) Proportional cellular frequencies in omental WAT, normalized by total cell number. (C) Gene ontology (GO) networks of significantly upregulated DEGs in insulin-sensitive WAT. (D) Gene ontology (GO) networks of significantly upregulated DEGs in insulin-resistant WAT. (E) Significantly altered GO terms in APC1 and (F) APC2 from insulin-sensitive and insulin-resistant WAT. (G) Feature plots displaying transcript levels of clock genes in insulin-sensitive and insulin-resistant WAT. Circled cell populations indicate APCs and adipocytes.

## DISCUSSION

Our study provides evidence that peripheral circadian impairment is a key element driving WAT fibrosis and a diminished response to IF. In wild-type animals, we demonstrated that IF improves metabolism and resolves age- and diet-associated adipose tissue fibrosis and metabolic dysfunction. While many studies have focused on the preventive effect of IF on obesogenic and diabetic progression, we aimed to test the therapeutic potential of IF on a more clinically relevant mouse model by integrating the adverse effects of age and chronic overnutrition (*i.e.,* high fat diet). As expected, the aged DIO animals exhibited much more severe metabolic impairment, allowing us to examine the magnitude of the therapeutic benefits of IF and whether it could restore impaired adipose tissue function. In alignment with previous studies, IF resulted in body weight and fat mass loss, reduction in hepatic lipid accumulation, and improved glucose and insulin homeostasis in aged DIO mice. However, in contrast to previous reports (Kim *et al*., 2017; Li *et al*., 2017; Martinez-Lopez *et al*., 2017), we observed metabolic improvements in the absence of adipose tissue thermogenesis. These earlier studies have reported WAT browning as a driving factor for IF- or calorie restriction (CR)-mediated metabolic benefits. However, aging has been shown to significantly impair the thermogenic capacity in mice due to the senescence of APCs (Berry *et al*., 2017). External stimuli, such as cold exposure, were not sufficient to induce *de novo* beige adipogenesis in aged mice, resulting in a failure to defend body temperature. Another study reported that depletion of healthy ILC2 in aged WAT instigate thermogenic failure in response to cold temperature (Goldberg et al., 2021). These findings demonstrated a largely diminished thermogenic response in unhealthy animals. Importantly, obese humans also display blunted BAT activity and reduced stimulation of energy expenditure in response to cold, further supporting the notion that thermogenic capacity is largely dampened in aged/obese subjects (Orava et al., 2013). Although we observed IF-induced metabolic improvements, we did not observe enhanced energy expenditure, which was aligned with the lack of thermogenic activation in the WAT. This suggests that WAT thermogenesis is not an indispensable factor in conferring IF-mediated benefits. Interestingly, several human studies have reported that diet-induced weight loss by alternate-day fasting or CR is independent of WAT browning or enhanced energy expenditure in both obese and non-obese humans (Barquissau et al., 2018; Trepanowski et al., 2017). This suggests that WAT thermogenesis is selectively observed in lean and healthy mice and may not entirely explain the beneficial effects of dietary interventions in different mouse models.

Here, we identify the anti-fibrotic effects within the APCs as a novel driver of IF-induced benefits. Notably, we identified the circadian regulation of APCs to promote healthy WAT remodeling, even in aged DIO mice. In mice that are lacking functional circadian clock (whole-body or APC-specific), these metabolic effects were largely or partially diminished. In contrast to our study, Chaix et al. reported that time-restricted feeding (TRF) exerts preventive effects against metabolic dysfunction in mice lacking a circadian clock (Chaix *et al*., 2019). However, they employed whole-body *Cry1*/*Cry2* KO, liver-specific *Bmal1* KO and liver-specific *Rev-erbα*/*β* KO mice to test the effects of TRF. Different clock mutant mice exhibit varying metabolic phenotypes and could lead to differential responses to dietary interventions. Additionally, while our study employed 2:1 IF, this study employed TRF. By restricting daily food intake to a certain timeframe (*e.g.,* 10-14 hrs/day feeding time combined with 10–14 hrs/day fasting time), TRF can mimic physiological circadian rhythms and impose rhythmicity in the peripheral metabolic organs, even in mice lacking a circadian clock, which can, in turn, drive diurnal rhythm in energy utilization and metabolic outputs (Pickel and Sung, 2020). Meanwhile, animals that are subjected to IF regimens such as 2:1 IF or alternate-day fasting that include prolonged fasting duration (*i.e.*, 24 hrs/day) do not impose a specific feeding time. Hence, the cycles of prolonged fasting may not be sufficient to drive significant metabolic improvements in the absence of restoration of functional rhythmicity. Nonetheless, we observed the upregulation of circadian pathways in APCs of IF-subjected aged DIO animals, suggesting that age- and diet-induced circadian dysfunction of WAT is capable of being restored. While our manuscript was being prepared, a study by Hepler *et al*. reported that TRF induces adipocyte thermogenesis through circadian regulation of creatine metabolism (Hepler *et al*., 2022). Similarly, our snRNA-seq data also showed evidence of upregulated thermogenic and lipid metabolism pathways in mature adipocytes of IF-treated WAT, demonstrating similar metabolic outcomes despite the difference in the circadian alignment of meal timing. This suggests that the cellular impacts of both regimens may be regulated by a common upstream pathway that can control divergent downstream pathways, including circadian rhythm, thermogenesis, and fibro-inflammation in various cell populations. Indeed, a recent study illustrated that exercise upregulates circadian rhythm gene expressions, such as *Dbp,* and reduces fibro-inflammatory phenotype of APCs (Yang et al., 2022). These data, in parallel with our results, collectively suggest that not only circadian-aligned cues (*e.g.,* TRF) but other physiological or nutritional cues, such as exercise and prolonged fasting periods, can enhance circadian rhythm regardless of their alignment with central or peripheral clock rhythmicity. In the current study, unlike our previous study, IF failed to increase energy expenditure in the aged DIO mice despite enhanced thermogenesis and lipid metabolism pathway in WAT. This may be associated with largely reduced thermogenic potential of aged DIO mice due to prolonged HFD feeding and ageing of mice with slightly reduced food intake (Berry *et al*., 2017; Starr and Saito, 2012). Additional future studies are warranted to identify the differences and similarities in the underlying mechanisms of these physiological adaptation processes to differing fasting regimens.

Interestingly, the snRNA-seq of human visceral fat revealed distinct gene enrichment within the adipocytes and APCs, depending on the patients’ insulin sensitivity status. While APCs from insulin-sensitive WAT displayed enrichment of genes associated with circadian rhythm and response to insulin, APCs from insulin-resistant WAT were enriched for inflammation, chemotaxis, and ECM-associated genes. These changes were observed in the absence of body weight differences, suggesting that metabolic health, particularly insulin sensitivity, could be influenced by a functional circadian clock in WAT, particularly APCs. Similarly, a study by Maury *et al*. reported that adipocytes and APCs from obese human omental fat demonstrated suppressed circadian clock genes due to NF-κB competing with *BMAL1* for the transcriptional regulation of downstream target genes (Maury *et al*., 2021). Another study by Gabriel *et al*. also reported disrupted circadian rhythmicity and mitochondrial metabolism in the skeletal muscle of individuals with type 2 diabetes, in the absence of difference in BMI (Gabriel et al., 2021). This suggests that metabolic health is largely controlled by functional circadian rhythm in peripheral organs, and that a cell- and tissue-specific dysfunctional circadian clock may contribute to various metabolic phenotypes.

Taken all together, our study provides new insights into the novel role of APC circadian rhythm in regulating WAT fibrosis, and thereby systemic glucose homeostasis and insulin sensitivity. Our results identify the peripheral circadian regulation of APCs as an indispensable mediator of IF-induced adipose remodeling and metabolic benefits.

## LIMITATION OF THE STUDY

We provide evidence that circadian rhythm of adipose progenitor cells (APCs) plays a pivotal role in stem cell plasticity and tissue remodeling. However, future research is warranted to address the following limitations of our study: 1) The mechanisms by which intermittent fasting (IF) reprograms APC circadian rhythm remains elusive. 2) Since we used male mice for our study, it is unclear whether our findings are consistent in female mice. However, our data from human fat tissue harvested from female patients show similar findings as mouse data. Thus, similar outcomes are expected from female mouse study. Nonetheless, female mice need to be used to precisely test the sexual dimorphism of IF effect. 3) Our study used a small number of human adipose tissues. To better support our conclusion, increased number of human tissues from both male and female patients will be highly suggested. 4) Lastly, constitutive deletion of *Bmal1* in PDGFRα-expressing cells may impact the development and function of non-adipose organs due to the wide expression pattern of PDGFRα. Thus, employing an inducible *Bmal1* knockout model will be ideal to study the role of circadian rhythm of APCs in the absence of adverse developmental consequences.

## METHODS AND PROCEDURES

### Animal experiments

All animal use protocols were approved by the Animal Care Committee of The Centre for Phenogenomics (TCP) conformed to the standards of the Canadian Council on Animal Care. All animals were housed in a specific pathogen-free facility in ventilated cages under controlled settings (22±1°C, with 30-60% humidity), 12hr light-dark cycle, and *ad libitum* access to water. C57BL/6J mice were bred from TCP in-house mouselines (#000664). For aged DIO studies, male mice at 8 weeks of age were fed with 45% fat diet (Research Diets, D12451) for 10 months prior to subjecting them to IF studies. *Bmal1* KO and APC-*Bmal1* KO animals were fed with normal chow (Harlan #2918; 17% fat).

### Genetic mouse models

Whole-body *Bmal1* heterozygous mice (*Bmal1* Het) were obtained from The Jackson Laboratory (#009100, C57BL/6J background). *Bmal1* KO mice were obtained by crossing *Bmal1* Het males and females. APC-specific *Bmal1* KO mice were generated by crossing PDGFRα-Cre (The Jackson Laboratory; #013148) males and *Bmal1* floxed (#007668) female mice. Both mouse lines were maintained on C57BL/6J background.

### Intermittent fasting

For each IF cohort, body weight-matched male mice were randomly divided into *ad libitum* (AL) and intermittent fasting (IF) groups. Mice were fed with either normal diet (ND) or 45% high-fat diet (HFD). Mice in the IF groups were subjected to 6 weeks of 2:1 IF regimen, which comprises of 1 day of fasting followed by 2 days of AL feeding (Kim *et al*., 2019). The food was removed and added at 12:00PM. Mice in the AL group were handled equally. Food intake and body weights were measured whenever food was added or removed.

### Body composition analysis

Body composition was analysed using EchoMRI-100 body composition analyzer (Echo Medical Systems, Texas, USA), which quantifies fat and lean mass in live, non-anesthetized mice.

### Energy expenditure analysis

Energy expenditure was evaluated through indirect calorimetry using Promethion Core systems (Sable Systems International, Las Vegas, USA). Mice were singly housed in Promethion control cabinets, which provides cage environment that matched the TCP facility. The system measures VO2, VCO2, food intake, water intake, locomotor activity, and respiratory quotient. The mice were provided with the same cage enrichment and were acclimated to the cages for 2 days prior to measuring their energy expenditure over the duration of one cycle of IF. The measurements were obtained every 15 minutes, which were averaged hourly for each mouse. To adjust for body weight differences between aged DIO-AL and IF mice, oxygen consumption rates of individual mice were plotted in relation to body mass and analyzed using ANCOVA. The plots were generated on https://calrapp.org.

### Glucose/insulin tolerance tests

For glucose and insulin tolerance tests (GTT and ITT, respectively), mice were fasted for 16 hours and 6 hours prior to the experiment, respectively. Mice were intraperitoneally injected with glucose (1mg/g of body weight) or insulin (Humulin, 0.65mU/g of body weight). Fasting glucose levels, and 5-, 15-, 30-, 60-, and 120-minutes of glucose levels post-glucose injection were measured using glucometer (Contour NEXT, Bayer Healthcare). Fasting glucose levels, and 15-, 30-, 60-, 90-, and 120-minutes of glucose levels post-insulin injection was measured. The homeostatic model assessment of insulin resistance (HOMA-IR) was calculated using the following formula: fasting glucose (μU/L) × plasma insulin (nmol/L) ÷ 22.5.

### Plasma insulin analysis

Mice were fasting for 24 hours prior to blood collection. Blood was collected in heparin- coated tubes and centrifuged at 5000rpm for 10 minutes at 4°C to separate the plasma. Plasma insulin level was quantified using an insulin ELISA kit (ALPCO Diagnostics, 80-INSMS-E01) according to the manufacturer’s instructions.

### Adipose tissue stromal cell isolation

For CyTOF and cell culture studies, stromal vascular fraction (SVF) was isolated from WAT. Visceral white adipose tissues are dissected, finely minced, and digested with Type II collagenase (Worthington; #LS004176) in a 37°C incubator for 30-40 minutes. Digested tissue was neutralized with DMEM/F12 media supplemented with 10% FBS and filtered using 100μm filter. The flow-through was centrifuged for 5 minutes at 400g. After removing the floating adipocyte fraction and supernatant, red blood cell lysis buffer (Sigma; #R7757) was added and incubated at room temperature for 2-3 minutes. The red blood cell lysis buffer is neutralized with DMEM/F12 media (supplemented with 10% FBS) and centrifuged for 5 minutes at 400g. The cell pellet was further processed for CyTOF or *in vitro* studies. For cell culture studies, the isolation process is conducted in a sterile BSC.

### CyTOF staining and data processing

The dissociated cells were washed in cell staining medium (CSM) and centrifuged for 5 minutes at 400g. Fc receptors are blocked by suspending the cell pellet in 25μL of 2X anti-mouse CD16/CD32 for 10-15 minutes at room temperature. Metal-tagged antibodies are prepared in 2X dilution in CSM. Antibodies used for CyTOF staining are listed in Table 2. 25μL of antibodies are added to the cells (without removing Fc receptor blocker) and are incubated at room temperature for 30 minutes. After washing out the antibodies with 2mL of CSM, the cells are pelleted by centrifuging for 5 minutes at 400g. After washing the cells with PBS, the cells are stained with 2X cisplatin viability stain for 5 minutes at room temperature and are consequently quenched with 2mL of CSM. The cells are washed with CSM, pelleted, and are stained with 1mL of Iridium intercalator diluted in PBS containing 1.6% PFA and 0.3% saponin. The cells are washed with CSM an hour before the acquisition. Final samples were suspended in Maxpar Cell Acquisition Solution containing EQ normalization beads (Fluidigm; #201237). Data is collected by Helios instrument. FCS files are normalized using EQ beads by the Helios software. Cytobank Enterprise (Beckman Coulter) was used to generate biaxial, histogram, and viSNE plots.

### In vitro adipogenic differentiation

After single cell dissociation PWATs from aged DIO-AL and -IF mice, the cells are seeded on 12- or 24-well plates, based on cell number in high-glucose DMEM (Gibco; #11965092), supplemented with 10% FBS, penicillin-streptomycin, gentamicin, sodium pyruvate and MEM non-essential amino acid solution. Once cells reach confluency, adipogenic differentiation medium (DMEM with 10% FBS, 0.5mM IBMX, 1μM dexamethasone, and 5μg/mL insulin). After 2 days, medium is changed to medium only containing insulin. Culture medium was changed every 2 days. On day 7-8 of culture, brightfield images were taken using Olympus CKX41.

### Treatment of 2D and 3D preadipocytes with Rev-erb**α** agonist SR9009 and immunofluorescent staining

Murine preadipocyte cell line 3T3-L1 cells (ZenBio) were seeded on 24-well plates (50,000 cells/well) and were grown with DMEM media (10% FBS) until 80-90% confluency was reached. SR9009 (Sigma-Aldrich, #554726) was dissolved in DMSO, and cells were treated with 0μM, 5μM, or 10μM of SR9009 for 24 hours. The cells were washed and were collected for gene expression analysis. For generation of 3D adipose tissue spheroids, 3T3-L1 cells and human visceral preadipocytes (Lonza Bioscience, PT-5005) were cultured in 2D until confluency. For human preadipocytes, cells were grown with Preadipocyte Growth Medium-2 BulletKit^TM^ (PGM-2^TM^, PT8002). Then, cells were seeded into either 96-well (50,000 cells/well) or 24-well (150,000 cells/well for 3T3-L1, 25,000/well for human preadipocytes) ultra-low attachment (ULA) plates (96-well ULA, Costar; 24-well ULA, Nunclon Sphera 3D Culture System, ThermoFisher) as previously described (Taylor *et al*., 2020). Treated 3D spheroids were assessed for collagen formation by immunostaining. Briefly, 3D spheroids were washed in PBS and blocked with PBS (5% goat serum and 0.3% Triton-X) for one hour. Then, 3D spheroids were incubated overnight with anti-collagen VI antibody (1:200, abcam, ab6588) at 4 °C. After washing in PBS, spheroids were incubated with secondary antibody for one hour in room temperature. After washing in PBS, the spheres were stained with DAPI for 3 minutes in room temperature. 3D spheroids were imaged by Nikon A1R confocal microscope.

### qPCR Analysis

Tissues or sorted cells were lysed using Trizol (ThermoFisher; #15596026). Total RNA was extracted from cells or tissues using RNeasy Mini Kit (Qiagen; #74104). Complementary DNA was synthesized from 200-2000μg of RNA using M-MLV reverse transcriptase (ThermoFisher; #28025-013). Gene expression assay was conducted using SYBR green PCR Mastermix (ThermoFisher; #4364346) on Quantstudio 5 (Applied Biosystems), and relative CT values are normalized with 36B4 gene. Primer sequences will be provided upon request.

### Histological Analysis

Freshly harvested tissues were fixed in 4% PFA and dehydrated in increasing concentrations of ethanol (70%, 95%, 100%). The tissues were incubated in xylene and wax for 3 hours each and were subsequently embedded in paraffin. Sections of 5μm were stained with hematoxylin and eosin, and random parts of the sections were selected for imaging. Images were obtained using Zeiss Axioscope. For adipocyte size analysis, CellProfiler (http://cellprofiler.org) was used to quantify adipocyte size. The pipeline to quantify adipocyte size in WAT was created by Lauren Pickel in Sung Lab. Images were converted to greyscale, uneven illumination was corrected, and noise was reduced before setting a threshold and recognizing objects. Extraneous objects were filtered out with parameters of solidity (minimum 0.75), area (500-13,000 pixels), and eccentricity (maximum 0.97). Adipocyte area was converted to square micrometers with a factor identified by scalebar measurement in ImageJ. For adipose tissue fibrosis and collagen staining, we performed histochemistry for Picrosirus Red staining in paraffin embedded tissue sections.

### Nuclei isolation

Approximately 50-70mg of frozen WAT samples were minced into smaller pieces on dry ice. Minced tissues were incubated in lysis buffer for 5 minutes. Supernatant and pellet were dounced, until the solution appeared milky. Isolated nuclei were washed with wash buffer and centrifuged at 800g for 10 minutes. The cells are filtered through 40μm cell strainer and stained with DAPI and sorted using FACS. After FACS, nuclei were stained with SYBR Green II (ThermoFisher; #S7654) and counted. Sorted nuclei were used as input (∼5,000 to 10,000 nuclei) into 10X Genomics single-cell 3’ v3.1 assay and processed according to the 10X Genomics protocol.

### Sequencing

The molarity of each library was measured by Bioanalyzer (Agilent Technologies) and qPCR amplification data, which was calculated based on the library size. Samples were pooled and normalized to 1.5nM. Library was denatured using 0.2N NaOH for 8 minutes at room temperature and was neutralized with 400mM Tris-HCl. Library pool was prepared at final concentration of 300pM and was loaded on Novaseq 6000 (Illumina). Samples were sequenced with the following run parameters: Read 1 - 28 cycles, Read 2 – 90 cycles, Index1 - 10 cycles, and Index2 – 10 cycles. Sequencing target read for each library was 50,000 reads per nuclei.

### Single nuclei data analysis

The 10X Genomics’ CellRanger v6.0.0 software was used for sequence alignment and barcode processing with mouse reference genome (mm10-2020-A). Removal of background (non-cellular) barcodes was initially performed using the default CellRanger knee-plot filtering, and potential doublets were removed using ‘scDblFinder’ (Germain et al., 2021). Low quality nuclei were also removed from the dataset by only retaining nuclei with more than 500 unique features (genes), less than 20000 total UMI counts (corresponding to ∼ 95^th^ percentile) and less than 1% mitochondrial transcripts. The filtered data was analysed using standard data integration protocol in Seurat v4.0.1, including log normalization, variable feature selection, selection of integration anchors, and Louvain clustering (Stuart et al., 2019). To identify cluster identities, our data was mapped on to reference dataset of white adipose tissue in mouse and humans as generated by Emont et al (Emont et al., 2022). Mapping of reference and our (target) datasets was conducted using Seurat’s integrated pipeline (as described in (Stuart *et al*., 2019), and the proportion of the nearest centroid in common reference space was used to determine putative cluster identities. Cell type annotations were also verified using cluster-specific genes as identified using the Wilcoxon Rank Sum test implemented in the FindAllMarkers function in Seurat. Next, differential gene expression between case and control samples was modelled using a hurdle MAST model (Finak et al., 2015). We defined genes with an absolute fold change of >0.2 and FDR corrected p-value of <0.05 to be differentially expressed between conditions. These differentially expressed genes were used for Gene Ontology (GO) enrichment analysis (Gene Ontology, 2001). GO analysis was conducted for biological processes ontology with minimum gene set size of 10 and maximum of 500, using clusterProfiler package (Yu et al., 2012). Module scores specific to biological processes (such as ‘circadian gene activity’) were calculated using Seurat’s ‘AddModuleScore’ function using genes from the mouse and human Gene Ontology annotations (for mouse and human snRNAseq datasets, respectively). The module score is the average expression of a set of genes (biological ‘program’) on the single cell level, subtracted by the aggregated expression of control feature sets calculated based on the binned average expression within each expression bin (Butler et al., 2018; Stuart *et al*., 2019). Lastly, to ascertain putative transcription factors that regulate differentially expressed genes between conditions, we used the ChEA 2016 transcriptional factors dataset (Lachmann *et al*., 2010). ChIP Enrichment Analysis (ChEA 2016) dataset includes published chromatin immunoprecipitation profiling studies from transcriptional factors in human and mouse cell lines, tissues, etc (Kuleshov *et al*., 2016; Lachmann *et al*., 2010). Specific to the focus of this study, we also added ChIP BMAL1 data from human omental fat tissue using the top 95^th^ percentile of BMAL1 regulated genes (Maury *et al*., 2021).

### Statistical Analysis

All results are presented as mean ± SEM. Statistical significance of differences between two groups was determined by two-tailed unpaired and paired Student’s t-test. Differences among the three or more groups were analyzed via one-way ANOVA followed by Sidak post-hoc test.

## ACKNOWLEDGMENTS

We thank Anthony Zhao from Dr. Cynthia Guidos’ laboratory and Tina Chen and Joe Cozzarin from the Center for Advanced Single Cell Analysis at the Hospital for Sick Children Research Institute for their considerable technical contribution, including optimization of staining and panel design in CyTOF experiments. We would like to additionally acknowledge Mandy Xu and Troy Ketela from Princess Margaret Genomics Centre for their technical contribution and guidance for single nuclei RNA sequencing experiments. For this research, H.-K.S. is supported by grants from Canadian Institute of Health Research (CIHR, PJT-162083), Natural Sciences and Engineering Research Council (NSERC, RGPIN-2016-06610) of Canada, and Sun Life Financial New Investigator Award of Banting & Best Diabetes Centre (BBDC) of University of Toronto. H.-K.S and J.-R. K. are supported by Korea-Canada Research Fund (2019K1A3A1A74107385) through the National Research Foundation of Korea (NRF) funded by the Ministry of Science and ICT. K.-H.K is supported by the National New Investigator Award from the Heart and Stroke Foundation of Canada. J.H.L. is supported by Doctoral Program Postgraduate Scholarship (PGS-D) from the Natural Sciences and Engineering Research Council (NSERC) of Canada.

## AUTHOR CONTRIBUTIONS

J.H.L and H.-K.S. conceived, designed, and performed the research. J.H.L, J.L-H.Y, Y.H.K, and K.H.K performed animal experiments, CyTOF, and RT-PCR. Y.P. performed analysis of single nuclei RNA sequencing datasets. L.P. provided histological analysis of animal tissues. K.E, and J.T, performed *in vitro* experiments. J-G.P analyzed bulk-tissue RNA sequencing datasets. T.J, A.O, and S.D harvested human adipose tissues and contributed to human fat tissue single nuclei RNA sequencing. J-R.K and S-Y.P supported single nuclei RNA and bulk-tissue RNA sequencing and discussed results. All authors discussed the results and agreed to the final version of the manuscript.

