## Supplementary Information for "Circadian reprogramming of adipose progenitor cells regulates intermittent fasting-mediated adipose tissue remodeling and metabolic improvement"

### SUPPLEMENTARY FIGURE LEGENDS

**Supplementary Figure 1. PWAT of aged DIO mice display pro-inflammatory and reduced adipogenic signatures.**

(A) Schematic diagram of young, DIO, aged, and aged DIO mouse model. (B) Body weights of young (n = 6), DIO (n = 6), aged (n = 5), and aged DIO (n = 6) mice. (C) IPGTT of four groups. (D H&E stained images of the liver. (E) Representative biaxial plots and bar graphs showing macrophage subtypes in young and aged DIO PWAT (n = 3 each). (F) Proportion of eosinophils in PWAT. (G) Proportion of ILC2 in PWAT. Values are presented as means ± SEM. *$p$<0.05, **$p$<0.01, ***$p$<0.005

**Supplementary Figure 2. IF reduces lipid droplet accumulation in the liver of aged DIO and improves insulin sensitivity.**

(A) Cumulative food intake over 6 weeks of IF, normalized by per gram of BW. (B) BAT and liver weight after 6 weeks of IF. (C) H&E stained images of BAT and liver. (D) Fasting insulin level after 24 hours of fasting. (E) HOMA-IR. Calculated by (fasting glucose × fasting insulin / 22.5). Values are presented as means ± SEM. *$p<0.05$, **$p<0.01$, ***$p<0.005$

**Supplementary Figure 3. IF does not enhance diet-induced thermogenesis in aged DIO mice.**

(A) Hourly oxygen consumption rate during one cycle of IF, normalized by body weight. (B) Oxygen consumption rate on fasting and feeding days. (C) Average oxygen consumption rate during one cycle of IF. (D) Linear regression analysis of oxygen consumption rate with body weight as a covariate during fasting and feeding days. (E) mRNA levels of thermogenic genes in PWAT. (F) mRNA levels of thermogenic genes in BAT. Values are presented as means ± SEM. *$p<0.05$, **$p<0.01$, ***$p<0.005$

**Supplementary Figure 4. snRNA-sequencing reveals immune and stromal cells that are altered by IF.**

(A) Top 3 differentially expressed genes of each cell cluster. (B) Transcript levels of various APC markers in two APC clusters. (C) GO term pathways significantly upregulated and downregulated in macrophage 1 cluster. (D) GO term pathways significantly upregulated and downregulated in macrophage 2 cluster. (E) Body weight during short-term IF. (F) Body composition after 3 weeks of IF. (G) Tissue weight after 3 weeks of IF. (H) Fasting glucose level after 3 weeks of IF. (I) GO term pathways significantly upregulated by short-term IF. Blue arrows indicate thermogenic and circadian pathways. (J) Top 20 differentially regulated genes by short-term IF.

**Supplementary Figure 5. Whole-body and APC-specific Bmal1 knockout mice display altered WAT phenotype.**

(A) Hourly oxygen consumption rate of *Bmal1* Ctrl and *Bmal1* KO mice. (B) Oxygen consumption rate during inactive (day) and active (night) phases. (C) Average oxygen consumption rate over a single cycle of IF. (D) Representative macroscopic image of WAT mass of *Bmal1* Ctrl and *Bmal1* KO mice. (E) Histogram showing CD9 expression in PDGFRα+ APCs in PWAT. (F) Transcript level of *Bmal1* in mature adipocytes, PDGFRα+ and PDGFRα- cells. (n = 3 for each group). (G) Transcript level of *Bmal1* and *Dbp* in in PWAT (n = 5-6 per group). (H) Hourly oxygen consumption rate of APC-*Bmal1* Ctrl and APC-*Bmal1* KO mice. (I) Oxygen consumption rate during day and night. (J) Average oxygen consumption rate. Values are presented as means ± SEM. *$p$<0.05, **$p$<0.01, ***$p$<0.005.

**Supplementary Figure 6. Bmal1 KO and APC-Bmal1KO mice display diminished metabolic benefits in response to IF.**

(A) Adipocyte size of IWAT. (B) Hourly oxygen consumption rate of *Bmal1* Ctrl-AL and *Bmal1* Ctrl-IF mice. (C) Average oxygen consumption rate. (D) Hourly oxygen consumption rate of *Bmal1* KO-AL and *Bmal1* KO-IF mice. (E) Average oxygen consumption rate. (F) Adipocyte size of IWAT. (G) Hourly oxygen consumption rate of APC-*Bmal1* Ctrl-AL and APC-*Bmal1* Ctrl-IF mice. (H) Average oxygen consumption rate. (I) Hourly oxygen consumption rate of APC-*Bmal1* KO-AL and APC-*Bmal1* KO-IF mice. (J) Average oxygen consumption rate. Values are presented as means ± SEM. *$p$<0.05, **$p$<0.01, ***$p$<0.005.

**Supplementary Figure 7. snRNA-seq reveals differential gene expressions in omental fat from insulin-sensitive and insulin-resistant individuals.**

(A) HOMA-IR in insulin-sensitive and insulin-resistant patients. (B) GO terms significantly upregulated and downregulated in adipocytes of insulin-sensitive individuals. (C) Schematic diagram displaying *in vitro* SR9009 treatment on 3T3-L1 cells. (D) Immunofluorescence staining of collagen type VI after treatment of SR9009.

### SUPPLEMENTARY TABLE LEGENDS

Supplementary Table 1. Characteristics and metabolic parameters of study subjects.

Supplementary Table 2. Antibody panel for adipose tissue mass cytometry analysis

Supplementary Figure 1.

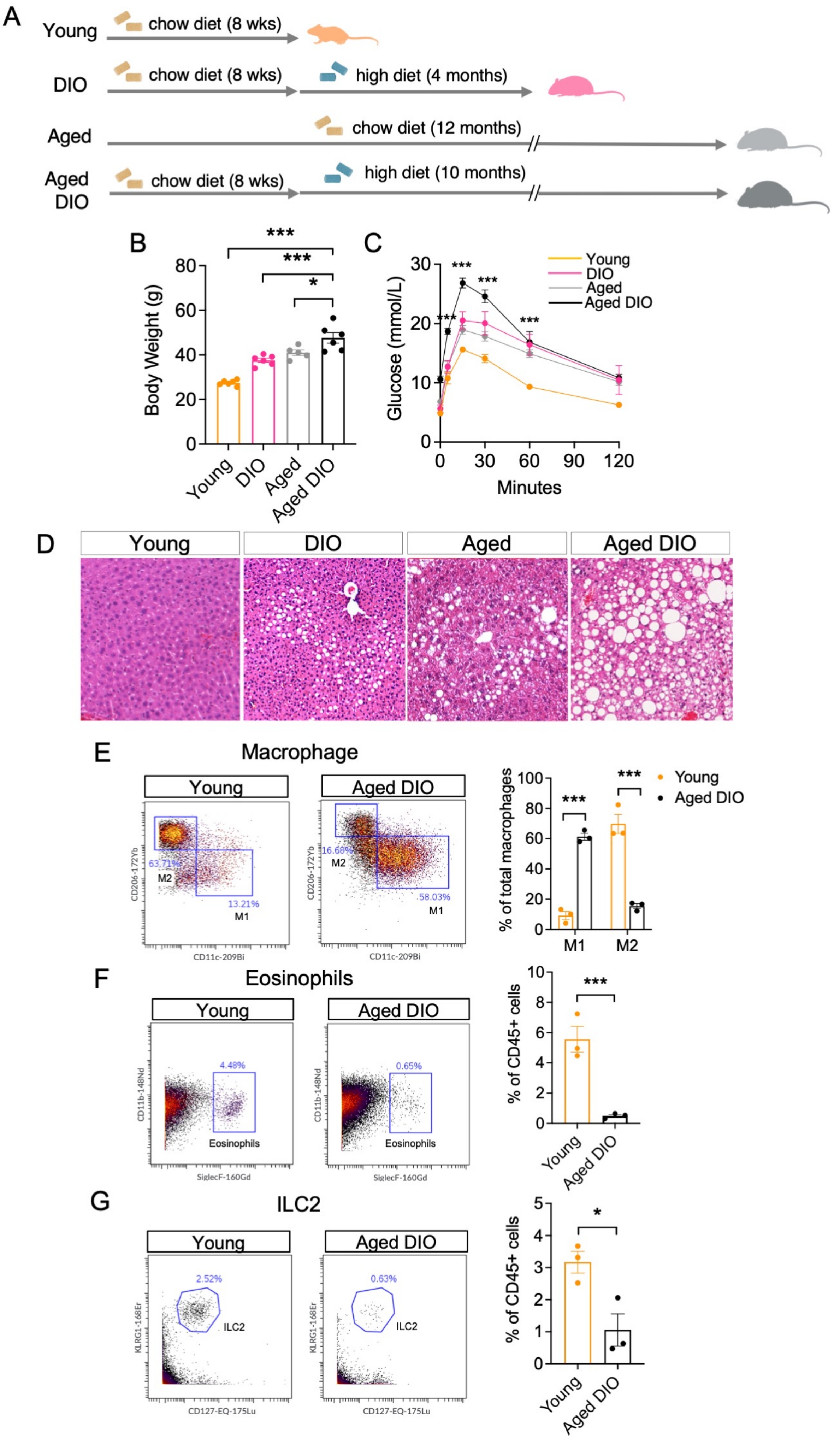

Supplementary Figure 2.

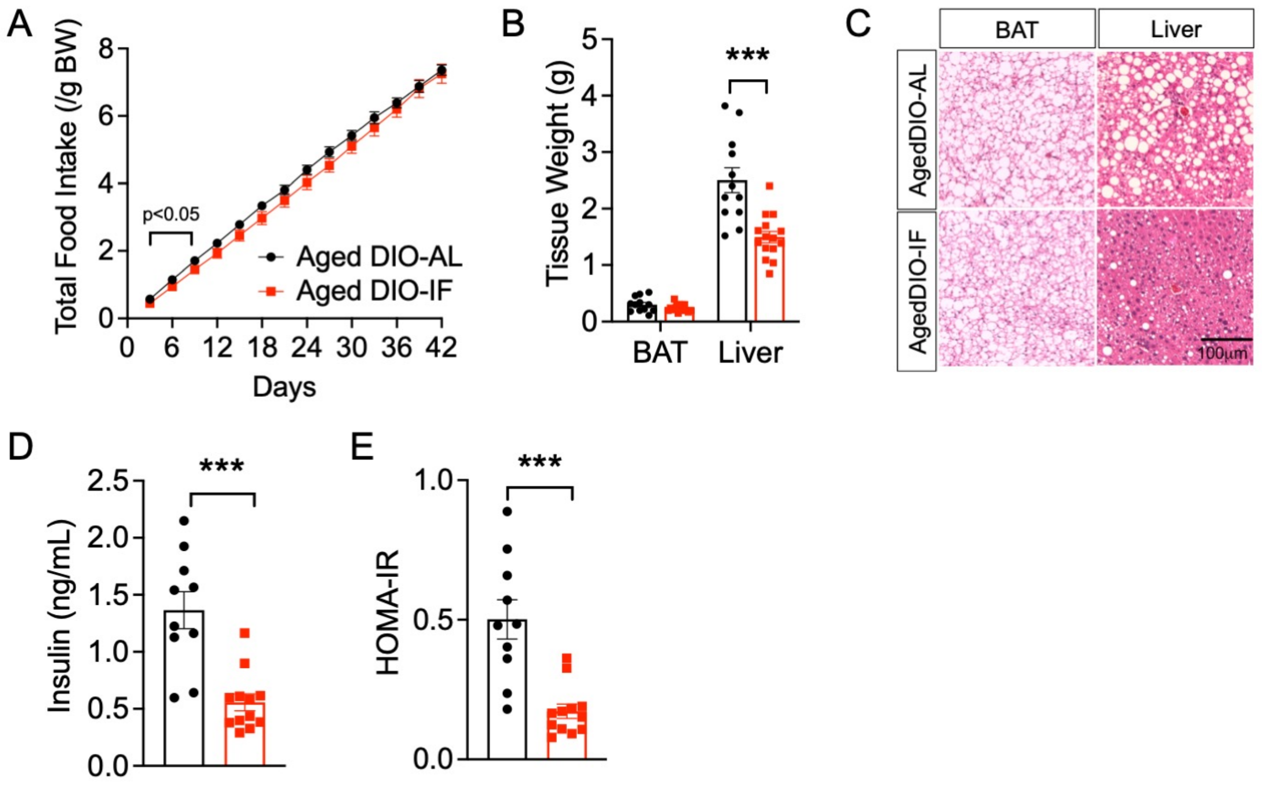

Supplementary Figure 3.

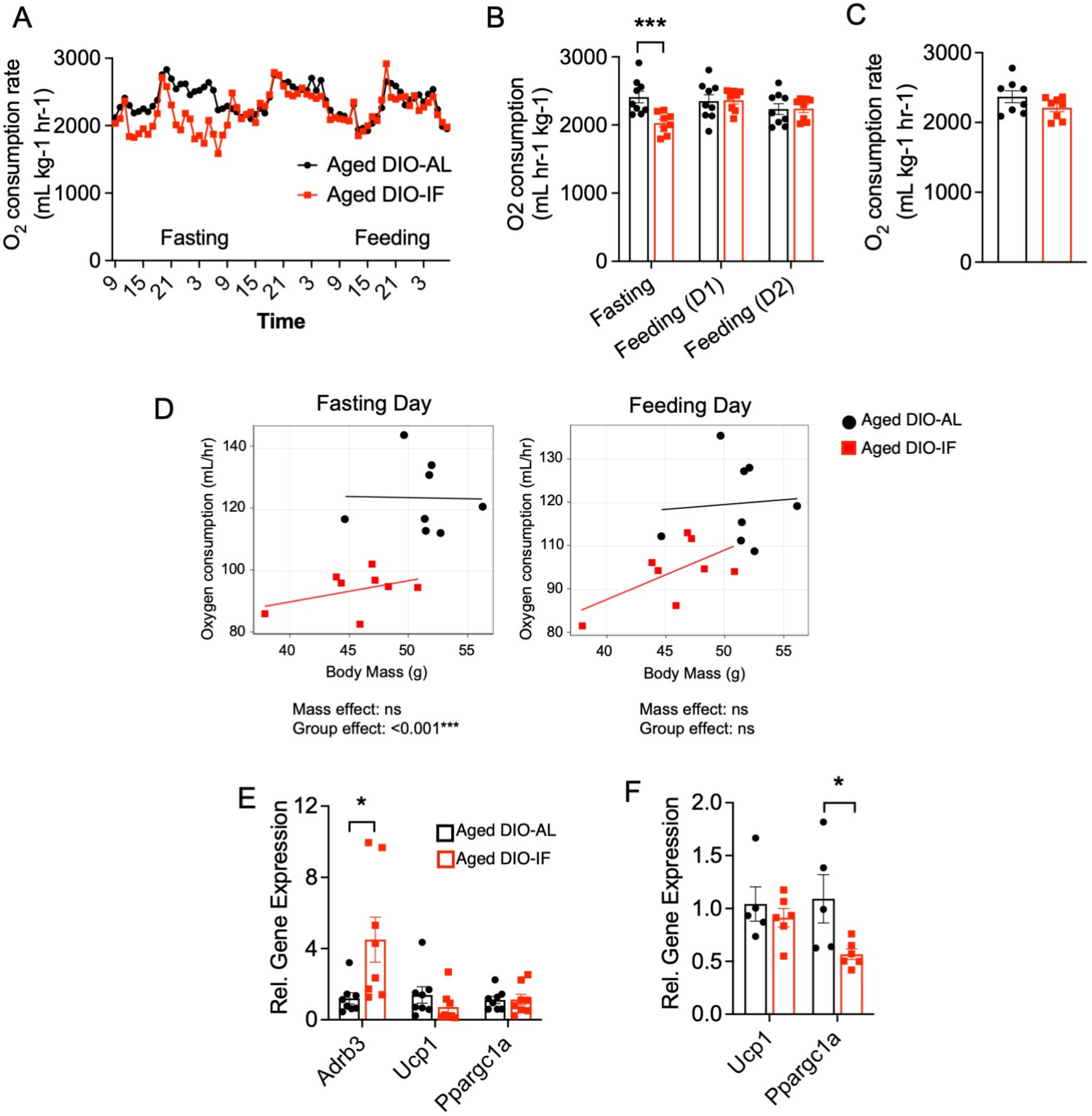

Supplementary Figure 4.

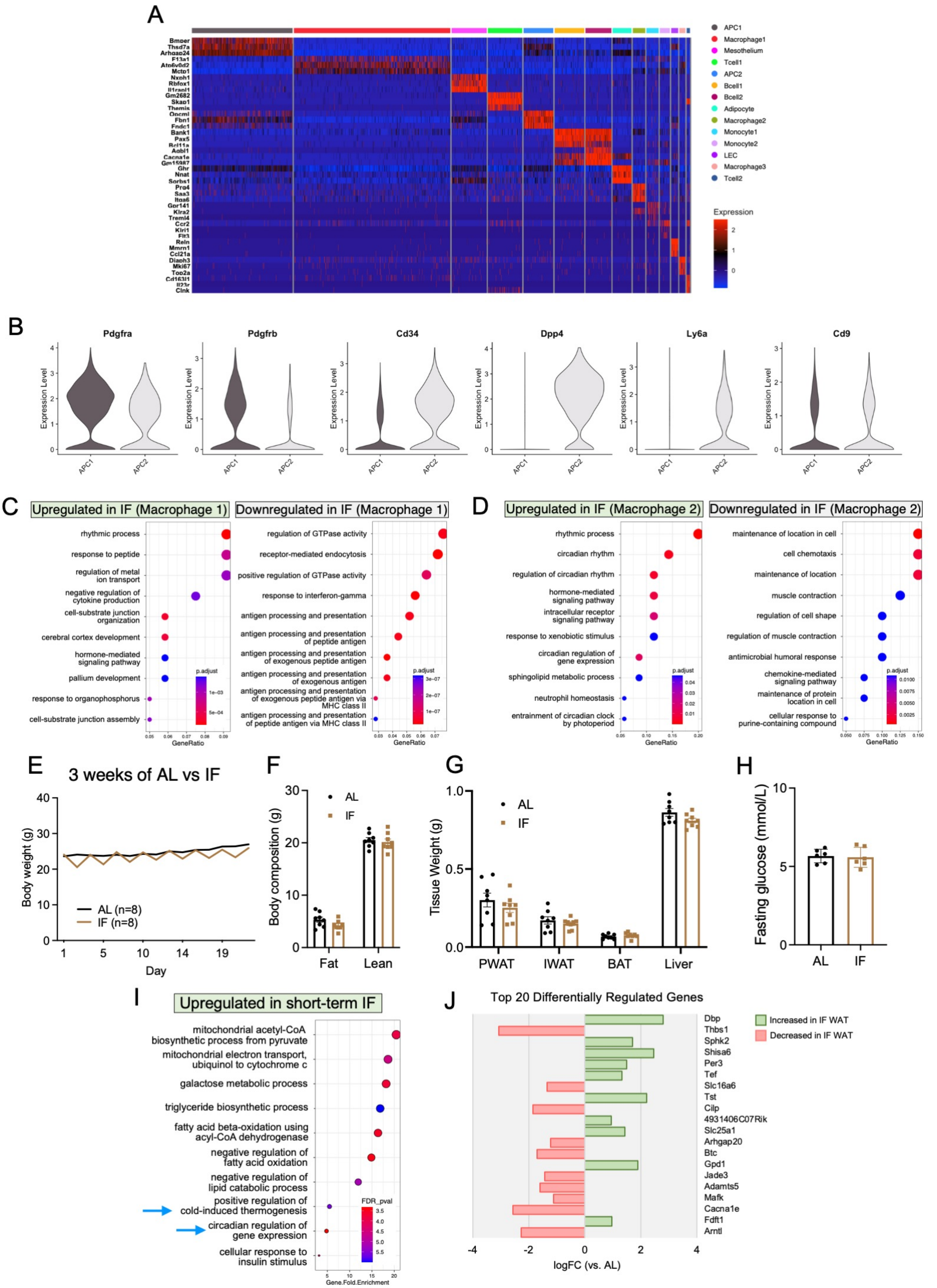

Supplementary Figure 5.

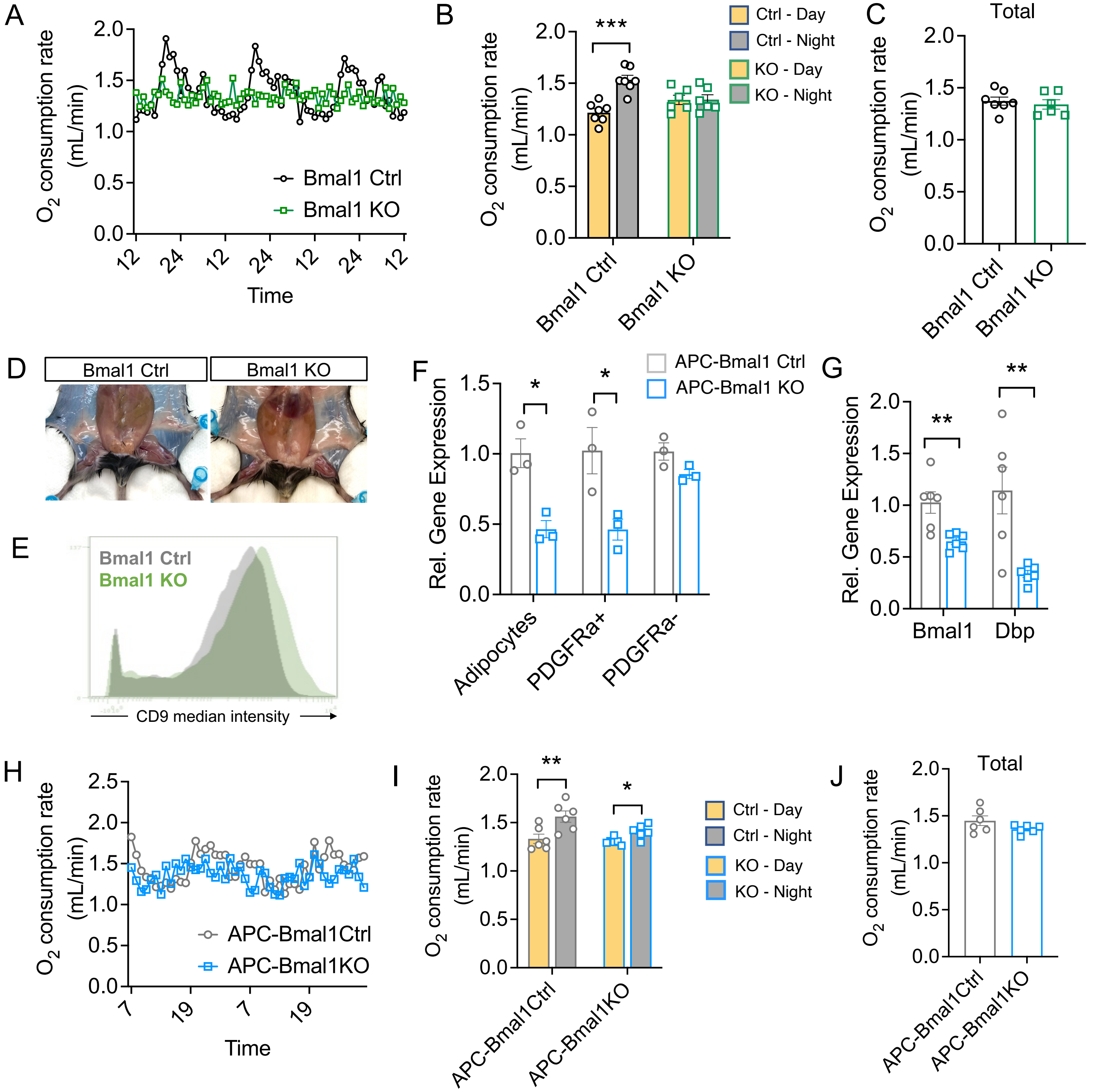

Supplementary Figure 6.

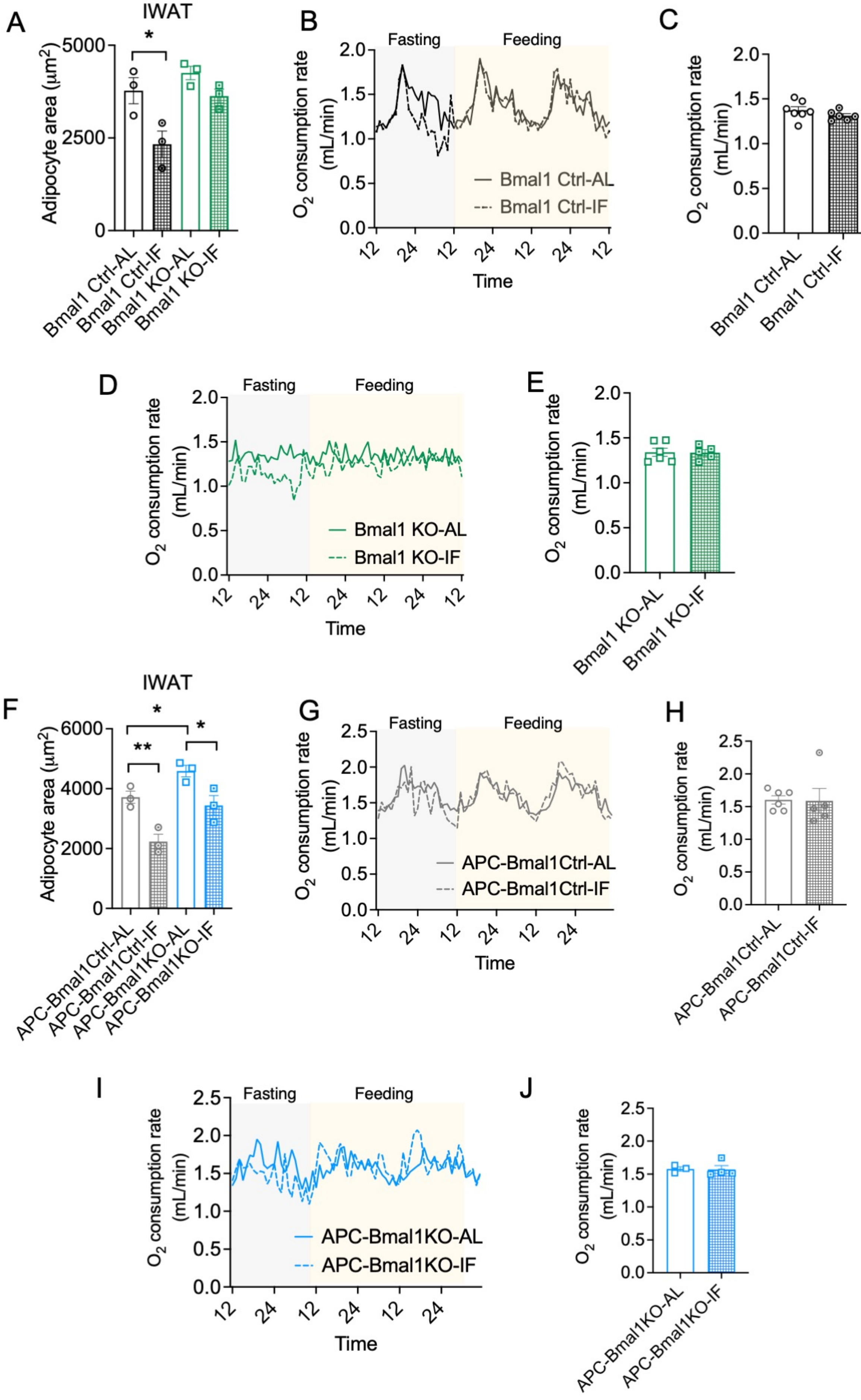

Supplementary Figure 7.

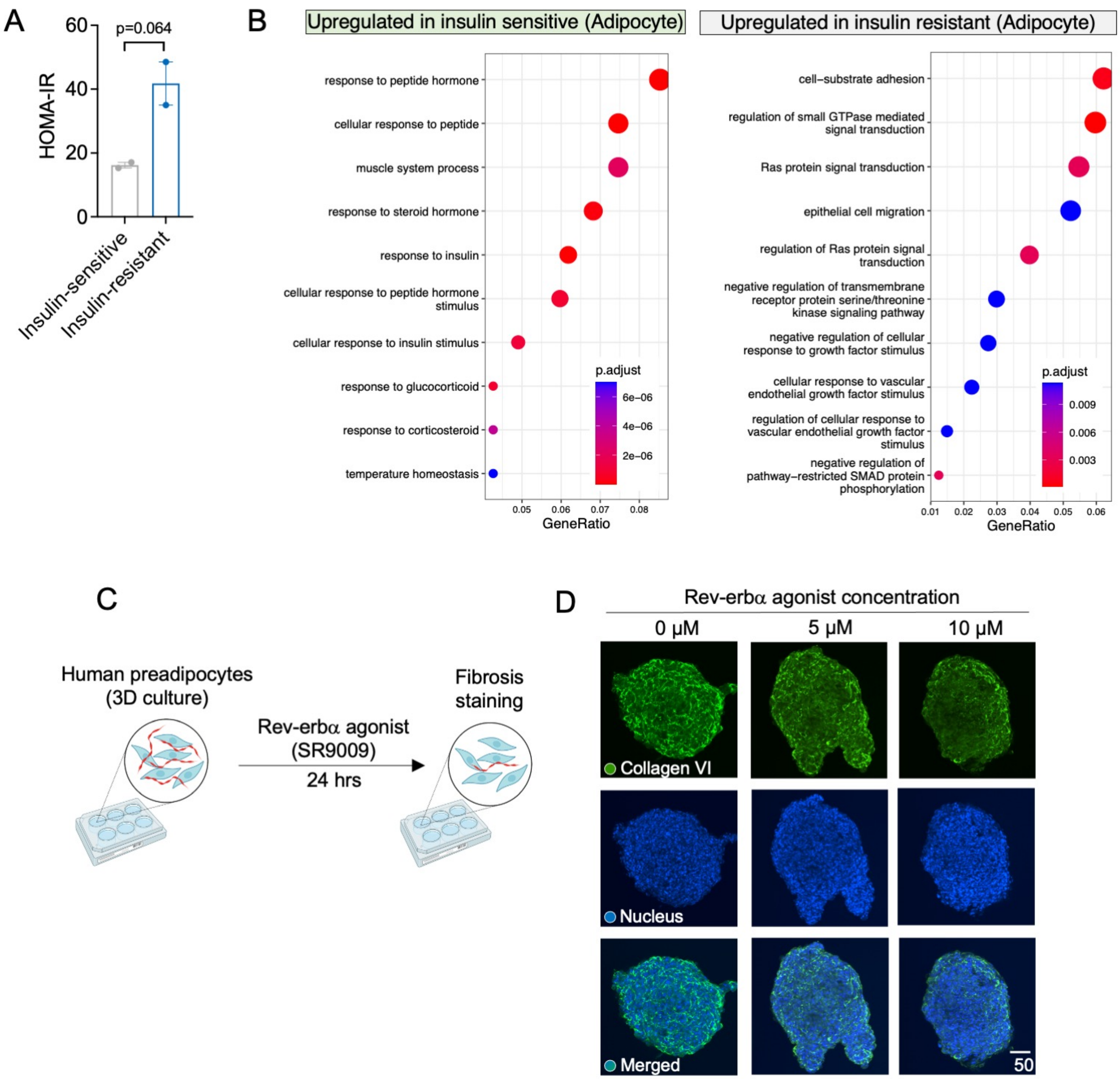

Supplementary Table 1. Characteristics and metabolic parameters of study subjects.

| | Age | Sex | Weight (kg) | Height (m) | BMI | Waist Circumference (cm) | Blood Pressure | Fasting Glucose (mmol/L) | Fasting Insulin | HOMA-IR | Triglyceride |
|---|---|---|---|---|---|---|---|---|---|---|---|
| S021 (Insulin-sensitive) | 63 | F | 173.5 | 1.637 | 64.7 | 162 | 137/87 | 4.8 | 80 | 2.82 | 1.48 |
| S049 (Insulin-sensitive) | 47 | F | 206.7 | 1.7 | 71.4 | 137 | 130/77 | 5.3 | 65 | 2.53 | 1.7 |
| S050 (Insulin-resistant) | 40 | F | 172 | 1.65 | 63.2 | 141 | 142/88 | 5.1 | 214 | 8.01 | 0.76 |
| S058 (Insulin-resistant) | 57 | F | 134.6 | 152.6 | 57.8 | 137.2 | n/a | 6.2 | 127 | 5.78 | 1.1 |

Supplementary Table 2. Antibody panel for adipose tissue mass cytometry analysis.

| **Antibody** | **Clone** | **Vendor** | **Concentration** |
|---|---|---|---|
| 89Y-CD45 | 30-F11 | Fluidigm | 1:300 |
| 115In-Ly6C | HK1.4 | Biolegend | 1:100 |
| 142Nd-CD11c | N418 | Biolegend | 1:300 |
| 143Nd-SiglecF | E50-2440 | BD | 1:1000 |
| 144Nd-MHC2 | M5/114.15.2 | Biolegend | 1:500 |
| 146Nd-PDGFRβ | APB5 | Biolegend | 1:100 |
| 148Nd-CD11b | M1/70 | Biolegend | 1:800 |
| 149Sm-CD19 | 1D3 | BD | 1:500 |
| 150Nd-CD24 | M1/69 (Maxpar ready) | Biolegend | 1:500 |
| 151Eu-CD31 | 390 (Maxpar ready) | Biolegend | 1:1000 |
| 152Sm-DPP4 | H194-112 | Biolegend | 1:800 |
| 154Sm-ICAM1 | YN1/1.7.4 | Biolegend | 1:1000 |
| 156Gd-PDGFRa | APA5 | ThermoFisher | 1:800 |
| 159Tb-CD64 | X54-5/7.1 | Biolegend | 1:300 |
| 160Gd-CD142 | AF3178 | R&D | 1:300 |
| 161Dy-CD25 | 3C7 | Biolegend | 1:100 |
| 163Dy-CD8b | H35-17.2 | ThermoFisher | 1:2500 |
| 164Dy-Sca1 | E13-161.7 | Biolegend | 1:1000 |

| | | | |
|---|---|---|---|
| 165Ho-CD4 | RM4-5 (Maxpar ready) | Biolegend | 1:300 |
| 166Er-CD9 | EM-04 | Novus | 1:500 |
| 167Er-Ly6G | 1A8 | Biolegend | 1:500 |
| 168Er-KLRG1 | 2F1 | BD | 1:1000 |
| 169Tm-TCRβ | H57-597 | Biolegend | 1:500 |
| 170ER-NK1.1 | PK136 | Biolegend | 1:100 |
| 171Yb-CD29 | HMβ1-1 | Biolegend | 1:1000 |
| 172Yb-CD206 | C068C2 | Biolegend | 1:800 |
| 173Yb-CD34 | RAM34 | ThermoFisher | 1:100 |
| 174Yb-CD3 | 145-2C11 (Maxpar ready) | Biolegend | 1:200 |
| 175Lu-CD127 | A7R34 (Maxpar ready) | Biolegend | 1:300 |
| 176Yb-B220 | RA3-6B2 | Biolegend | 1:500 |
